# Tissue- and ethnicity-independent hypervariable DNA methylation states show evidence of establishment in the early human embryo

**DOI:** 10.1101/2021.12.17.473110

**Authors:** Maria Derakhshan, Noah J. Kessler, Miho Ishida, Charalambos Demetriou, Nicolas Brucato, Gudrun E. Moore, Caroline H.D. Fall, Giriraj R. Chandak, Francois-Xavier Ricaut, Andrew M. Prentice, Garrett Hellenthal, Matt J. Silver

## Abstract

We analysed DNA methylation data from 30 datasets comprising 3,474 individuals, 19 tissues and 8 ethnicities at CpGs covered by the Illumina450K array. We identified 4,143 hypervariable CpGs (“hvCpGs”) with methylation in the top 5% most variable sites across multiple tissues and ethnicities. hvCpG methylation was influenced but not determined by genetic variation, and was not linked to probe reliability, epigenetic drift, age, sex or cell heterogeneity effects. hvCpG methylation tended to covary across tissues derived from different germ-layers and hvCpGs were enriched for associations with periconceptional environment, proximity to ERV1 and ERVK retrovirus elements and parent-of-origin-specific methylation. They also showed distinctive methylation signatures in monozygotic twins. Together, these properties position hvCpGs as strong candidates for studying how stochastic and/or environmentally influenced DNA methylation states which are established in the early embryo and maintained stably thereafter can influence life-long health and disease.

## Introduction

DNA methylation (DNAm) plays a critical role in mammalian development, underpinning X-chromosome inactivation, genomic imprinting, silencing of repetitive regions and cell differentiation^1^. DNAm states that vary between individuals have been a focus of Epigenome-Wide Association Studies (EWAS) due to their potential to drive phenotypic variation^2,3^. Factors influencing interindividual methylation differences include genetic variation^4,5^, cell heterogeneity effects^6,7^, sex^8,9^, age^10,11^, and pre- and post-natal environment^12–14^. Growing evidence from studies investigating DNAm patterns in multiple tissues suggests that these factors have both shared and tissue-specific influences on DNAm variation^12,15–18^.

In this study, we sought to identify loci with high interindividual methylation variability in multiple tissues and ethnicities, and to gain insights into the biological mechanisms influencing methylation variation. By using a large number of diverse sample types, we reasoned that identified loci would be robust to tissue-specific drivers of methylation variability such as those mentioned above, and to dataset-specific technical artefacts, including batch effects and poorly performing probes^19–22^. We began by characterising hypervariable CpGs (‘hvCpGs’) covered on the widely used Illumina HumanMethylation450K (hereafter ‘Illumina450K’) array^23^ that showed high interindividual variability across multiple datasets covering 19 different tissue/cell types and 8 ethnicities spanning a wide range of ages. We next investigated the influence of genetic variation, sex, age and probe reliability on methylation variability at hvCpGs. We additionally determined whether methylation states at hvCpGs covary across tissues by exploring their overlap with loci at which methylation varies between individuals but is correlated across tissues within a given individual, termed systemic interindividual variation or ‘SIV’. Since loci showing SIV have been linked to variable methylation establishment before germ-layer differentiation^24–27^, we further explored evidence for early embryo methylation at hvCpGs by determining their overlap with loci that show unique methylation patterns in MZ twins^25,28^ and with loci that show sensitivity to the periconceptional environment^29^. We assessed the genomic context of hvCpGs by exploring their association with multi-tissue histone marks and their proximity to transposable elements and regions of parent-of-origin-specific methylation. Finally, we probed putative functional roles of hvCpGs by interrogating EWAS trait associations and by performing gene ontology enrichment analysis.

Our curated list of hvCpGs show evidence of establishment in the early embryo and of correlation across tissues. They therefore serve as a useful resource for studying the influence of early environmental and/or stochastic effects on DNAm in diverse tissues and ethnicities, and for studying the impact of DNAm differences on life-long health and disease.

## Materials and Methods

### Methylation data used for identifying hvCpGs

Publicly available methylation data were downloaded from The Cancer Genome Atlas (TCGA) (https://www.cancer.gov/tcga) and the Gene Expression Omnibus (GEO)^30^ databases as methylation Beta matrices (Supplementary Tables 1 and 2). TCGA methylation data were downloaded using the *TCGAbiolinks (v2*.*18*.*0)* R package^31–33^, selecting only samples annotated as ‘Solid Tissue Normal’. Of the 33 TCGA datasets, 10 were selected for our study as these had methylation data in at least 20 samples. GEO methylation Beta matrices were downloaded from 11 unique accessions using the *GEOquery (v2*.*58*.*0)* R package^34^. Where available, detection p-values (measuring signal intensity), and metadata on age, sex, and disease status were also downloaded. We split GEO beta matrices into separate groups based on ethnicity and tissue/cell type and refer to the resulting 17 separated groups as ‘datasets’. Non-public datasets internal to this study include IlluminaEPIC^35^ array data from whole blood samples from Gambian 8-9-year olds (ISRCTN14266771^36^) and Illumina450K data from Bornean and Kenyan saliva samples^37^ (Supplementary Table 3). For IlluminaEPIC datasets we selected probes covered on the Illumina450K array. In total, we analysed 30 datasets (3 internal, 10 TCGA and 17 GEO) that covered 8 ethnicities and 19 different tissue/cell types (Supplementary Table 4).

### Methylation data processing

For each methylation dataset used in our main analysis, we used the *ChAMP (v2*.*20*.*1)* R package^38^ to remove: i) probes with a detection p-value > 0.01 in > 5% samples (where detection p-values were available), ii) probes mapping to multiple genomic positions^39^, iii) probes mapping to the X and Y chromosomes, and iv) single nucleotide polymorphism (SNP)-related probes identified by Zhou *et al*.^39^ that contain SNPs (MAF > 1%) within 5 bp of the CpG interrogation site and/or SNPs effecting probe hybridisation. Where ethnicity information was available, we removed probes with population-specific SNPs identified by Zhou *et al*. using 1000 Genomes populations (MAF > 1%), otherwise we removed the General Recommended Probes^40^. Probes that had a missing value in any of the samples in a specific dataset were removed from that dataset. To reduce technical biases introduced by differing type I and type II probe designs on the Illumina450K and IlluminaEPIC arrays, we applied Beta Mixture Quantile normalisation (BMIQ)^41^ to all datasets using the *champ*.*norm()* function from the *ChAMP* R package. All datasets were adjusted for the first 10 principal components (PCs) of variation to account for methylation variability driven by known and/or unknown technical artefacts (such as plate and array position) and cell heterogeneity. Methylation values were adjusted for these 10 PCs, age (where available) and sex by taking the residuals from a linear regression on methylation M values, where M is defined as *log*_*2*_*(beta/1-beta)*. Finally, for each probe, we removed outlier methylation values, defined according to Tukey’s outer fences (Q1 – 3*IQR and Q3 + 3*IQR). The hg19 reference genome was used throughout all relevant analyses as the Illumina450K array metadata manifest uses this version.

### Identification of hvCpGs

For each dataset, we identified CpGs within the top i% of CpGs by methylation Beta variance in ≥ j% of datasets in which the CpG was covered, for increasing values of *i* and *j*. We additionally required that selected CpGs were covered in a minimum of 15 datasets (Supplementary Fig. 1A). To define the ethnicity- and tissue-independent hypervariable CpGs (hvCpGs) explored in this paper, we set the threshold at *i, j* = 5,65 (Supplementary Fig. 1B).

### Probe reliability

Technically unreliable probes were identified by examining intra-class correlation coefficients (ICCs) from two studies. The first study compared methylation consistency between the Illumina450K and IlluminaEPIC platforms using 365 blood DNA samples, defining poor quality probes as those with ICC ≤ 0.4^22^. The second study examined methylation reliability between technical replicates from 265 African American peripheral blood leukocyte samples on the Illumina450K platform, defining poor quality probes as those with ICC ≤ 0.37^42^. We defined technically unreliable probes as those reported as being poor quality in at least one of these two studies (Supplementary Table 5).

### Methylation quantitative trait locus (mQTL) analysis

mQTL summary statistics from the Genetics of DNA Methylation Consortium (GoDMC), a meta-GWAS of 36 European blood cohorts (N = 27,750) generated using imputed genotype data (∼10 million SNPs) and ∼420,000 CpGs^43^ were used for this analysis. Significance thresholds of p < 1×10^−8^ and p < 1× 10^−14^ were applied for *cis* and *trans* mQTLs respectively^43^, giving 271,724 significant SNP-CpG associations comprising 190,102 CpGs and 224,648 SNPs. The variance in DNA methylation explained by a given mQTL was estimated as 2 * β * MAF(1-MAF), where β is the effect size and MAF is the minor allele frequency^44^.

### Monozygotic twin discordance

We analysed CpGs identified as being ‘equivalently variable’ between MZ co-twins and between unrelated individuals (‘evCpGs (blood)’) by Planterose Jiménez *et al*.^45^ using Illumina450K data in whole blood. 154 of these evCpGs replicated in adipose tissue from 97 MZ twin pairs (‘evCpGs (blood & adipose)’). evCpGs are candidates for methylation states that are established stochastically after MZ twin splitting.

### Control CpG sets

#### Distribution-matched controls

hvCpGs are enriched for intermediate methylation states (Supplementary Fig. 2). This property of hvCpGs has the potential to bias several downstream analyses, for example because this can affect power to find association with phenotypes in EWAS. We therefore constructed a set of CpGs with similar distribution of methylation Beta values to hvCpGs in the Caucasian blood dataset (‘Blood_Cauc’, Supplementary Table 1). This dataset was chosen as it has the highest number of post-natal samples and because several downstream analyses leverage published studies that used blood methylation data. For each of the 4,108 hvCpGs covered in the ‘Blood_Cauc’ dataset, a two-sided Kolmogorov-Smirnov (KS) test (*ks*.*test()* in R) was used to test for the divergence in methylation Beta distributions between the hvCpG and technically reliable (see ‘Probe reliability’, Methods) background probes, selecting the CpG with the greatest p-value (requiring a p-value > 0.1). In total, 3,566 hvCpGs were each matched to a control CpG (‘distribution-matched controls’, Table 1, Supplementary Fig. 3).

**Table 1.** Main CpG sets used in this study.

| CpG set | n | Notes |
| --- | --- | --- |
| hvCpGs | 4143 | CpGs within top 5% methylation Beta variance in at least 65% datasets in which the CpG is covered, requiring the hvCpG to be covered in at least 15 datasets and to be reported as technically reliable. |
| array background | 406306 | CpGs covered in at least 15 of the 30 datasets used in this study. |
| distribution-matched controls | 3566 | Array background CpGs with similar methylation Beta distributions to hvCpGs in the 'Blood_Cauc' dataset, requiring each control CpG to be technically reliable. |
| de-clustered hvCpGs | 2640 | A set of hvCpGs in which no CpGs is within 4kb of another CpG. |
| mQTL-matched controls | 3722 | CpGs reported by the GoDMC meta-GWAS<sup>41</sup> with the same number of mQTL associations and similar mean % variance explained by an mQTL, requiring each control CpG to be present in at least as many datasets as the hvCpG. |

#### mQTL-matched controls

To determine the degree to which hypervariability at hvCpGs is explained by mQTL effects, each hvCpG was matched to a CpG amongst those reported in the GoDMC meta-analysis^43^. Controls were selected to have i) the same number of mQTL associations, ii) a similar mean % variance explained by mQTL (across all significant mQTL) and iii) presence in at least as many datasets as the hvCpG (Table 1, Supplementary Fig. 4).

### Identification of hvCpG clusters

hvCpG clusters were identified by considering the decay of methylation correlation with distance at hvCpGs. To do this, we calculated the average pairwise Spearman correlation (*ρ*) across hvCpG pairs with inter-CpG distance falling within 100 bp bins, for datasets with at least 100 samples (Supplementary Fig. 5B). The distance threshold for defining hvCpG clusters was chosen to be 4,000 bp as this is approximately the point at which pairwise correlations levelled out (Supplementary Fig. 5B). In total, 2,219 (54%) hvCpGs fell into 716 clusters comprising at least 2 CpGs, with the remaining 1,924 (46%) hvCpGs falling outside of these clusters (Supplementary Fig. 5C). In 563 (79%) of these clusters, the average Spearman correlation (*ρ*) across hvCpG pairs was > 0.5 (Supplementary Fig. 5D).

### ‘De-clustering’ of hvCpGs

To account for the possibility that our analyses may be biased by the non-random distribution and inter-dependence of hvCpGs in CpG clusters, we generated a de-clustered set of hvCpGs in which no CpG was within 4 kb of another CpG. 2,640 de-clustered hvCpGs were generated by randomly selecting one CpG from each of the clusters and then including all ‘singleton’ CpGs falling outside of clusters.

### Age stability

To examine temporal stability of hvCpGs we used published intra-class correlation coefficients (ICCs) for probes on the Illumina450K array determined using white blood cell samples taken ∼6 years apart^46^. The ICC scores compare within-sample variability (across the two time-points) to between-sample variability, with ICC ≥ 0.5 defined as temporally stable by Flanagan *et al*.^46^ Because methylation data from the Flanagan *et al*. dataset were publicly available (GSE61151), we compared ICC scores at hvCpGs to those at CpGs with similar methylation Beta distributions to hvCpGs at the first time point (Supplementary Fig. 6A) to ensure that high hvCpG ICC scores were not biased by the high variability of hvCpGs. These CpGs were matched to each hvCpG using the same Kolmogorov-Smirnov method detailed in ‘Distribution-matched controls’ but using the Flanagan *et al*. methylation data instead the ‘Blood_Cauc’ dataset^44^. Longer-term susceptibility to epigenetic drift was examined by determining the proportion of hvCpGs that overlap a published set of 6,108 CpGs identified using whole blood Illumina450K data from 3,295 individuals aged 18 to 88 years that show an increased methylation variability with age of more than 5% every 10 years^11^ (Supplementary Fig. 6B).

### Published CpG sets used to investigate early embryo establishment

We used the following publicly available data to examine evidence that methylation states at hvCpGs are established in the early embryo. See Table 2 for a summary of these datasets.

**Table 2.** Published CpG sets used in this study.

| CpG set | Description | n. CpGs overlapping array background | Reference |
| --- | --- | --- | --- |
| SIV | Interindividual methylation variation with concordant methylation across tissues derived from different germ layers within a given individual. See Supplementary Table 8. | 3089 | Harris *et al.*, van Baak *et al.*, Kessler *et al.*, Gunasekara *et al.* <sup>24-27</sup> |
| ESS | Greater-than-expected methylation similarity between MZ co-twins | 1217 | van Baak *et al.* <sup>25</sup> |
| MZ twinning CpGs | Probes differentially methylated between MZ and DZ twins. | 728 | van Dongen *et al.* <sup>28</sup> |
| evCpGs | MZ co-twin methylation discordance that is equivalent to methylation discordance between unrelated individuals in whole blood. A subset of these replicated in adipose tissue. | 317 (blood)<br>145 (blood & adipose) | Planterose Jiménez *et al.* <sup>45</sup> |
| SoC | CpGs at which methylation is associated with season of conception in Gambian children. | 242 | Silver *et al.* <sup>29</sup> |
| PofOm | Regions of parent-of-origin-specific methylation identified in peripheral blood from Icelandic individuals. | 732 CpGs in 116 PofOm regions | Zink *et al.* <sup>68</sup> |

SIV = systemic interindividual variation, ESS = epigenetic supersimilarity, evCpGs = equivalently variable CpGs, SoC = season-of-conception, PofOm = parent-of-origin-specific methylation.

#### Systemic Interindividual Variation (‘SIV’) CpGs

SIV-CpGs were collated from four published datasets that used either whole genome bisulfite sequencing (WGBS) or Illumina450K data from multiple tissues derived from different germ layers to identify CpGs displaying high interindividual variation and low intra-individual (cross-tissue) variation. These properties are suggestive of variable methylation establishment before germ layer differentiation^24–27^. Further details on the four SIV screens used in this study are given in Supplementary Table 6.

#### Epigenetic Supersimilarity (‘ESS’) CpGs

Epigenetic supersimilarity (ESS) loci were identified by van Baak *et al*.^25^ using Illumina450K data from adipose tissue from 97 MZ and 162 dizygotic (DZ) twin pairs^47^. In that study, 1,580 ESS sites were identified within the top decile of methylation variance, with an interindividual methylation range > 0.4 and greater-than-expected concordance in MZ twins vs DZ twins. This supersimilarity is attributed to methylation establishment before MZ twin splitting.

#### MZ twinning CpGs

Van Dongen *et al*.^48^ performed an epigenome-wide association analysis on each of 6 cohorts with methylation data from both MZ and DZ twins (5 blood and 1 buccal) to identify probes differentially methylated between MZ twins and DZ (dizygotic) twins. A meta-analysis was then performed using the blood datasets to identify 834 Bonferroni-significant differentially methylated CpGs, which we refer to as ‘MZ twinning CpGs’.

#### Season of conception (‘SoC’) CpGs

Silver *et al*.^29^ used Illumina450K data to identify 259 CpGs associated with season-of-conception (‘SoC’) in Gambian 2-year olds with a minimum methylation difference of 4% between the peaks of the Gambian rainy and seasons.

#### Transposable elements and telomeres

Locations of ERV1 and ERVK transposable elements determined by RepeatMasker were downloaded from the UCSC annotations repository as previously described^26^. Telomere coordinates were downloaded from the UCSC hg19 annotations repository. (http://genome.ucsc.edu).

#### Imprinted genes, parent-of-origin-specific methylation (PofOm)

Imprinted genes classified as ‘predicted’ or ‘known’ were downloaded from https://www.geneimprint.com. Parent-of-origin-specific CpGs were identified by Zink *et al*.^49^ using WGBS data from peripheral blood from Icelandic individuals.

### SIV power calculation

To assess power to detect SIV in previous screens with small numbers of samples, we analysed the 4-individual multi-tissue dataset used by van Baak *et al*.^25,50^. We downloaded this dataset from GEO (GSE50192), selecting the same tissues (gall bladder, abdominal aorta sciatic nerve) used by van Baak *et al*.^25^. For each of the 1,042 SIV-CpGs reported by van Baak *et al*.^25^, we generated methylation values for three tissues for each simulated individual by randomly sampling from a 3-dimensional multivariate normal distribution, with mean equal to the mean of each tissue’s sampled methylation values at the CpG, and standard deviation specified by a 3×3 cross-tissue co-variance matrix of the sampled methylation values at the CpG. For each SIV-CpG, we sampled four simulated individuals and determined if this random sample met the SIV definition specified by van Baak *et al*.^25^, repeating this process 1000 times to give a power estimate (Supplementary Fig. 7).

### Processing and analysis of fetal multi-tissue dataset

The fetal multi-tissue dataset comprised 60 samples, corresponding to 30 individuals each with methylation data from two tissues derived from different germ layers (ectoderm: brain, spinal cord, skin; mesoderm: kidney, rib, heart, tongue; endoderm: intestine, gut, lung, liver). These fetal tissues were obtained from the ‘Moore Fetal Cohort’ from the termination of pregnancies at Queen Charlotte’s and Chelsea Hospital (London, UK). Ethical approval for obtaining fetal tissues was granted by the Research Ethics Committee of the Hammersmith, Queen Charlotte’s and Chelsea and Acton Hospitals (2001/6028). DNA was extracted from fetal tissues using the AllPrep DNA/RNA/Protein Mini Kit (Qiagen) and bisulfite conversion was carried out using EZ DNA Methylation Kits (Zymo Research). Samples were then processed using the Illumina InfiniumEPIC array. Derived methylation data were imported as .*idat* files into R and analysed using the *meffil* R package (*v 1*.*1*.*2*)^51^ with default parameters. Briefly, methylation predicted sex was used to remove 2 sex outliers (samples with methylation > 5 SDs from mean). Next, 1 sample was removed for which the predicted median methylation signal was more than 3 SDs from the expected signal, leaving 57 samples. 515 probes with detection-p-value value > 0.1 and 307 probes with bead number < 3 in more than 20% of samples respectively were removed. Array data were then corrected for dye-bias and background effects and functional normalisation was applied, specifying the number of PCs to be 7 (the PC at which the variance explained at control probes levelled out). Next, the *ChAMP (v2*.*20*.*1)* R package^38^ was used to remove cross-hybridising and multi-mapping probes, probes on XY chromosomes, and SNP-related probes, leaving 746,492 CpGs. We selected the 452,016 probes that overlapped the Illumina450K array and the 27 individuals for which both tissue samples passed quality control: 9 individuals with methylation data from endoderm and mesoderm, 10 individuals with methylation data from endoderm and ectoderm and 8 individuals with methylation data from mesoderm and ectoderm (see Supplementary Table 7). Methylation was then adjusted for predicted sex and batch using a linear model. For the 9 individuals with available endoderm-mesoderm samples we calculated the Pearson r between germ layer methylation values for each hvCpG, and repeated this for individuals with endoderm-ectoderm and mesoderm-ectoderm samples. The inter-germ layer correlation was then defined as the average Pearson r across these three comparisons. Following van Baak *et al*.^25^, interindividual variation was determined by calculating the mean methylation value across both tissues within each of the 27 individuals, before taking the range of these means for every CpG.

### Chromatin states at hvCpGs

Chromatin states were predicted by a ChromHMM 15-state model^52^ using Chromatin Immunoprecipitation Sequencing (ChIP-Seq) data generated by the Roadmap Epigenomics Consortium^53^. These data were downloaded for H1 ESCs (E003), fetal brain (E071), fetal muscle (E090), fetal small intestine (E085), foreskin fibroblasts (E055), adipose (E063) and primary mononuclear cells (E062) from the Washington University Roadmap repository. Chromatin states were collapsed into 8 states for clarity (Supplementary Table 8).

### EWAS trait associations at hvCpGs

hvCpG trait associations were determined using the EWAS catalogue (http://ewascatalog.org/), which details significant results (p-value < 1 × 10^−4^) from published EWAS studies. Considering only those traits for which at least 1% of hvCpGs overlapped associated CpGs (highlighted in green in Supplementary Table 9), we first extracted the array background CpGs overlapping the ‘Blood_Cauc’ dataset that were associated with each trait. We then calculated the proportion of these CpGs that comprised hvCpGs and blood distribution-matched controls (Table 1). Traits that were significantly enriched or depleted for hvCpGs relative to controls were those for which bootstrapped 95% confidence intervals did not overlap.

### Gene ontology term enrichment analysis

Gene Ontology (GO) term enrichment analysis was performed using the *missMethyl* R package (*v1*.*24*.*0*)^54^ using the *gometh()* function, setting arguments sig.cpg = hvCpGs, all.cpg = array.background, sig.genes = T, collection = “GO”, array.type = “450K” and prior.prob = T to adjust for variation in the number of 450K probes mapping to each gene.

### Bootstrapped confidence intervals

All bootstrapped 95% confidence intervals were calculated over 1,000 bootstrap samples.

## Results

### Identification of hypervariable CpGs

We analysed methylation data from 3,474 individuals across 30 datasets (28 Illumina450K and 2 EPIC array) comprising 19 unique tissue/cell types and 8 ethnicities covering a range of ages (Supplementary Tables 1-4). We focussed on CpGs covered by the Illumina450K array and began by excluding probes with poor detection p-values, cross-hybridising probes, probes on the X and Y chromosomes and probes associated with known SNPs (see Methods for details).

We aimed to identify CpGs with consistently high interindividual variation in methylation across diverse datasets, so minimising the effects of dataset-specific drivers of variability. We reasoned that removal of unmeasured technical, batch and cell heterogeneity effects within each dataset would maximise power to detect true biological variation across datasets and therefore adjusted all methylation values for the first ten principal components (PCs) of methylation variation, as well as sex (in datasets with both sexes) and age (where available).

Our strategy for identifying tissue- and ethnicity-independent hypervariable CpGs (‘hvCpGs’) is summarised in Fig. 1 and detailed in ‘Methods’. We first selected CpGs within the top *x*% of each dataset by methylation Beta variance, and then took the intersection of these CpGs across an increasing proportion of covered datasets, ensuring that each CpG was present in at least 15 datasets (Supplementary Fig. 1A). Using this approach, we identified 4,330 hvCpGs, defined as CpGs with methylation Beta variance in the top 5% of all CpGs in at least 65% of datasets for which that CpG passed QC criteria (Table 1). This definition met our required criteria of selecting CpGs that are highly variable in a large number of tissues and ethnicities (median [IQR] for each hvCpG = 13[10,15] and 7[6,7] respectively; Supplementary Fig. 1B). Note that no CpGs are expected to meet these criteria if the top 5% most variable CpGs in each dataset are entirely independent of those in the others. While re-defining these thresholds will change the set of hvCpGs, we noted that ∼80% of identified hvCpGs overlapped with an alternative set obtained when selecting CpGs with methylation Beta variance in the top 20% in at least 90% of datasets (Supplementary Fig. 1C), meaning that the majority of hvCpGs are within the top 20% of variable loci in almost all covered datasets.

**Figure 1.**
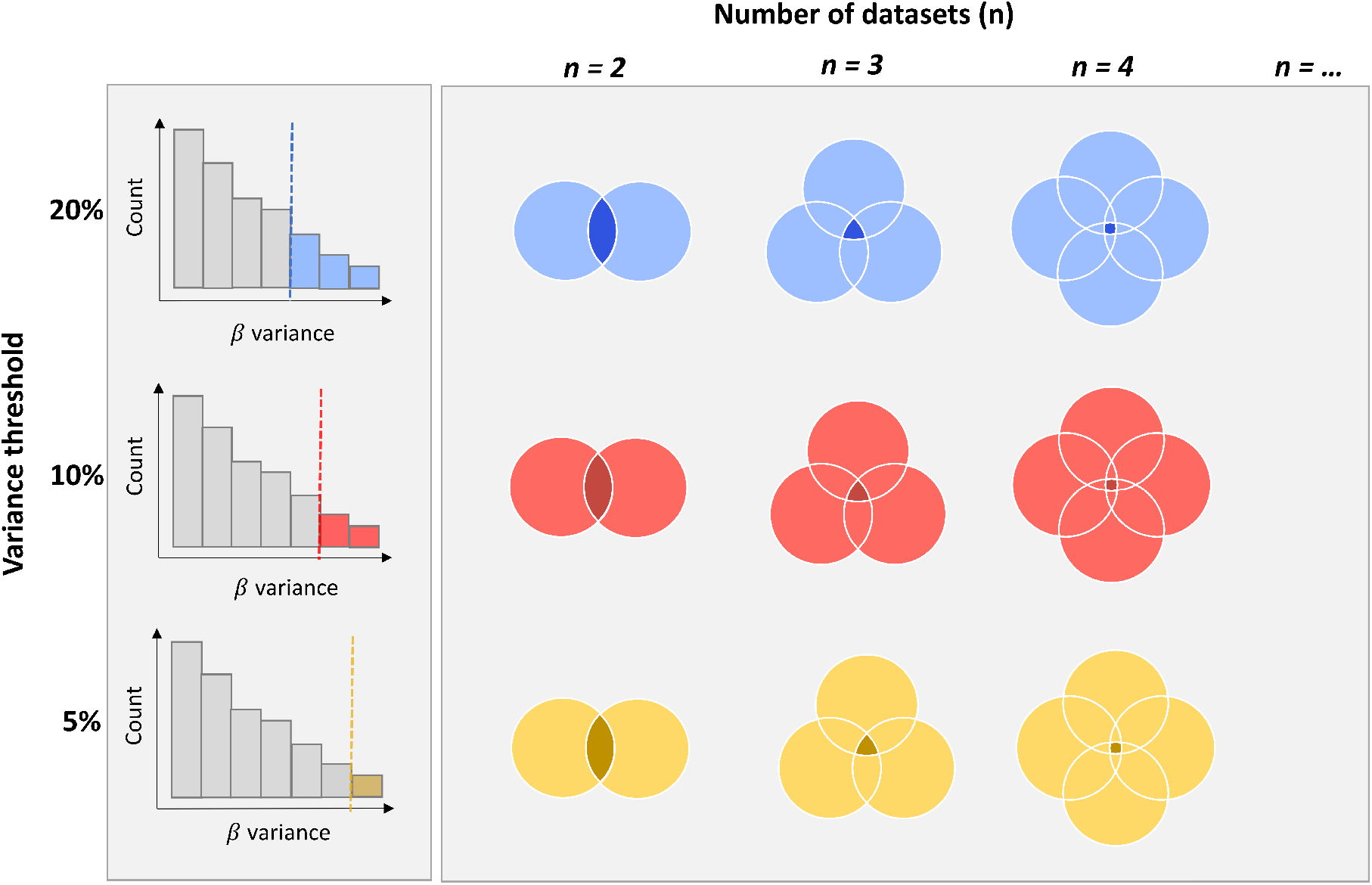
Schematic of the method for identifying tissue- and ethnicity-independent hypervariable CpGs (hvCpGs). The top 20%, 19%, 18% … 1% of variable CpGs by methylation Beta variance were first extracted from each of the 30 methylation datasets used in this study. The intersection of these CpGs was then taken over an increasing number of datasets (*n*), requiring each CpG to be present in a minimum of 15 out of the 30 datasets analysed (Supplementary Fig. 1).

We next compared the 4,330 hvCpGs with an alternative set obtained using the same method but without prior adjustment of each dataset for the first ten PCs. This alternative set contained only 1,302 CpGs, which confirmed our intuition that PC adjustment maximises power to identify true dataset-independent hypervariability by removing unwanted technical variation (Supplementary Fig. 8). Finally, we used reported measures of methylation variability among technical replicates^22,42^ to remove 187 technically unreliable probes (see Methods), leaving a final set of 4,143 hvCpGs (Table 1; Supplementary Table 5).

hvCpGs are enriched for intermediate methylation values in all datasets compared to the array background (Supplementary Fig. 2; see Table 1 for definition of array background) and are distributed throughout the genome (Supplementary Fig. 5A), with 2,219 (54%) falling within 716 ‘clusters’ containing two or more hvCpGs separated by < 4 kb (Supplementary Fig. 5C). To account for the possibility that our downstream analyses may be biased by these distributional properties, we generated a set of controls that were distribution-matched in a whole blood dataset (Supplementary Fig. 3) and a set of ‘de-clustered hvCpGs’ (Table 1, ‘Methods’).

### hvCpG variability is not driven by age, sex, or cell heterogeneity

Evidence from multiple studies suggests that methylation variability can increase with age (termed epigenetic drift)^11,55^, raising the possibility that cross-dataset hypervariability of hvCpGs is driven in part by a large proportion of adult/elderly samples. However, 3,815 (92%) out of 4,122 hvCpGs with methylation measured in cord blood and/or buccal samples from infants showed methylation variance within the top 5% of CpGs in those datasets (Supplementary Table 10), suggesting that high variability at hvCpGs arises in early life. We further probed age stability of hvCpGs by leveraging two studies of age effects in blood. The first study reported methylation consistency in individuals sampled at two time points six years apart^46^. The temporal stability of hvCpGs was significantly greater than that of controls with similar methylation Beta distributions to hvCpGs at the first time point (Wilcox paired signed-rank test p-value < 5.7 ×10^−81^), with 95% of hvCpGs considered temporally stable versus 89% of controls (Supplementary Fig. 6A). The second measured epigenetic drift in a cross-sectional study of 3,295 whole blood samples from individuals aged 18 to 88^11^. Only 7% of hvCpGs overlapped CpGs that show increased methylation variability with age, compared to 16.5% of blood distribution-matched controls (Supplementary Fig. 6B). This suggests that that the majority of hvCpGs are stable over a broad time period in whole blood and further supports the notion that hypervariability of hvCpGs in multiple datasets is not an artefact of epigenetic drift effects.

While methylation values were pre-adjusted for sex in all datasets where sex was available as a covariate (24 out of 30 datasets), we further investigated the potential for sex effects to drive methylation variance by considering the four female-only datasets. 4,102 (99%) of the 4,136 hvCpGs covered in any of these datasets had methylation variance among the highest 5% in at least one (Supplementary Table 10). Furthermore, we found no significant difference in mean methylation at hvCpGs between male and females in a diverse set of tissues (Supplementary Fig. 6C). Finally, 3,548 (96%) of the 3,678 hvCpGs covered in purified CD4+ and CD8+ datasets had methylation variance among the top 5% in at least one dataset (Supplementary Table 10). Together, these data strongly suggest that variability at hvCpGs is not driven by sex, age, or cell heterogeneity effects.

### Hypervariability is not driven by genetic variants

Genetic variation is an important driver of interindividual methylation differences^4,5^. There is evidence that mQTLs can be shared across different tissues^15,16,56,57^ and ethnic groups^5^, raising the possibility that ‘universal’ (multi-tissue and multi-ethnic) mQTLs might drive cross-dataset variability at hvCpGs. We therefore investigated the potential influence of methylation quantitative trait loci (mQTL) on methylation variability at hvCpGs by leveraging a recent large meta-GWAS (36 cohorts, n = 27,750 individuals) that identified common genetic variants associated with methylation in blood from Europeans^43^, reasoning that by definition ‘universal’ mQTLs would be included in this meta-analysis.

We considered multiple methylation variance thresholds (5%, 10% and 20%) and observed a positive relationship between hypervariability and both the probability of a significant mQTL association and the mean mQTL effect size (Fig. 2A). Amongst hvCpGs, there were 6,985 *cis* mQTL (covering 3,635 hvCpGs and 6,417 SNPs) and 971 *trans* mQTL (covering 713 hvCpGs and 753 SNPs). Overall, 3,722 (90%) of hvCpGs were reported to have at least one (*cis* or *trans*) mQTL association that were estimated to explain, on average, 4% of methylation variance (Fig. 2B). This suggests that additive genetic effects explain a small to moderate proportion of methylation variability at these hypervariable loci in blood. Noting that the statistical power to detect mQTL associations will be greater at loci that are inherently variable, we matched hvCpGs to CpGs with the same number of mQTL associations and similar average % variance explained by mQTL (‘mQTL-matched controls’, Table 1, Supplementary Fig. 4). hvCpGs showed an average 5-fold increase in methylation variance compared to mQTL-matched controls across datasets (Fig. 2C), further supporting the notion that methylation variation at hvCpGs is not principally driven by universal genetic effects.

**Figure 2.**
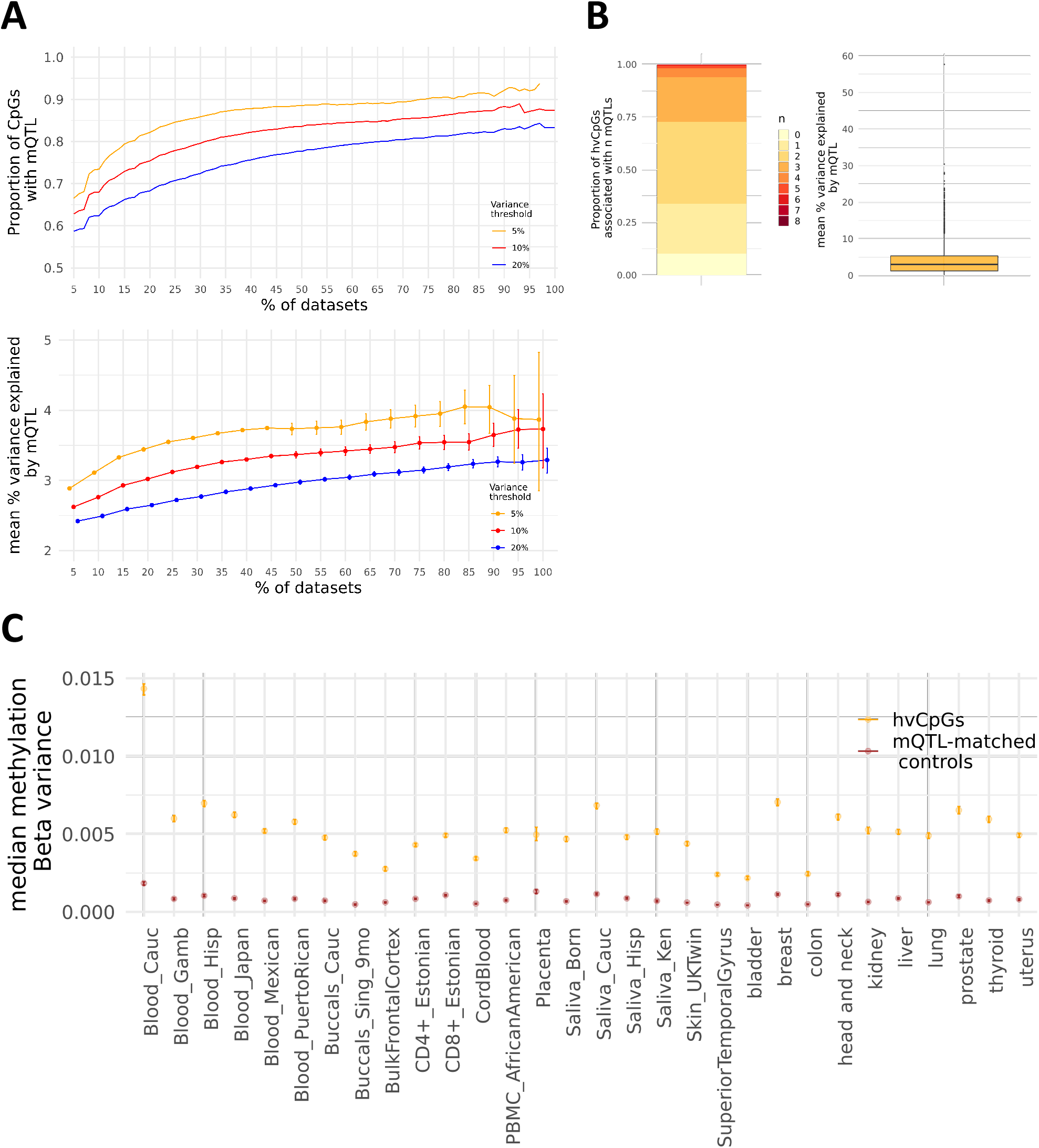
Genetic effects at hvCpGs using mQTL data from a large meta GWAS in blood (Min et al. 2021). **A)** The relationship between hypervariability and the proportion of CpGs with at least one mQTL association (top) and the mean mQTL effect size (bottom). Coloured curves represent CpGs with top 5% (orange), 10% (red) and 20% (blue) methylation Beta variance in at least *x%* of datasets. **B)** The distributions of the number of mQTL associations (left) and mean % variance explained by mQTL (right) at hvCpGs. **C)** Median methylation Beta variance at 3,722 hvCpGs overlapping the ‘Blood_Cauc’ dataset (orange) and corresponding controls matched on number of mQTL associations and mean % explained by mQTL (‘mQTL-matched controls’, Table1; Supplementary Fig. 4), in each dataset. Error bars in A and C are bootstrapped 95% confidence intervals. Note, error bars in C are very small.

To further probe the influence of genetic effects on hvCpG methylation we examined the overlap between hvCpGs and CpGs that show DNAm variation between monozygotic (MZ) co-twins that is *equivalently variable (ev)* to that between unrelated individuals, suggestive of genetically independent variable methylation establishment after MZ twin splitting^45^. In total, hvCpGs comprise 122 (42%) of the 317 evCpGs identified in blood (1.9-fold enrichment relative to distribution-matched controls) and 62% of those that were replicated as evCpGs in adipose tissue (2.8-fold enrichment relative to controls) (Supplementary Table 11), supporting the notion that hvCpGs are likely influenced but not determined by genetic variation in multiple tissues.

### hvCpGs show covariation across tissues derived from different germ layers

Variable DNAm states that covary across tissues derived from different germ layers and that are influenced but not determined by genotype may have been established before germ layer separation in early embryonic development^26^. None of the datasets considered here had multi-tissue data from the same individuals. We therefore examined the overlap between hvCpGs and 3,089 CpGs that show systemic (cross-tissue) interindividual variation (SIV), collated from four published sources^24–27^ (Supplementary Table 6). 24% of hvCpGs overlap a known SIV-CpG as do 21% of hvCpGs with a blood distribution-matched control, the latter representing a ∼5-fold enrichment (Fig. 3A, Supplementary Table 12, Supplementary Fig. 10A). We note that a further 540 (13%) hvCpGs are within 1 kb of a SIV-CpG, ∼5-fold greater than array background CpGs. This suggests that many hvCpGs directly overlap or co-localise with a known SIV-CpG.

**Figure 3.**
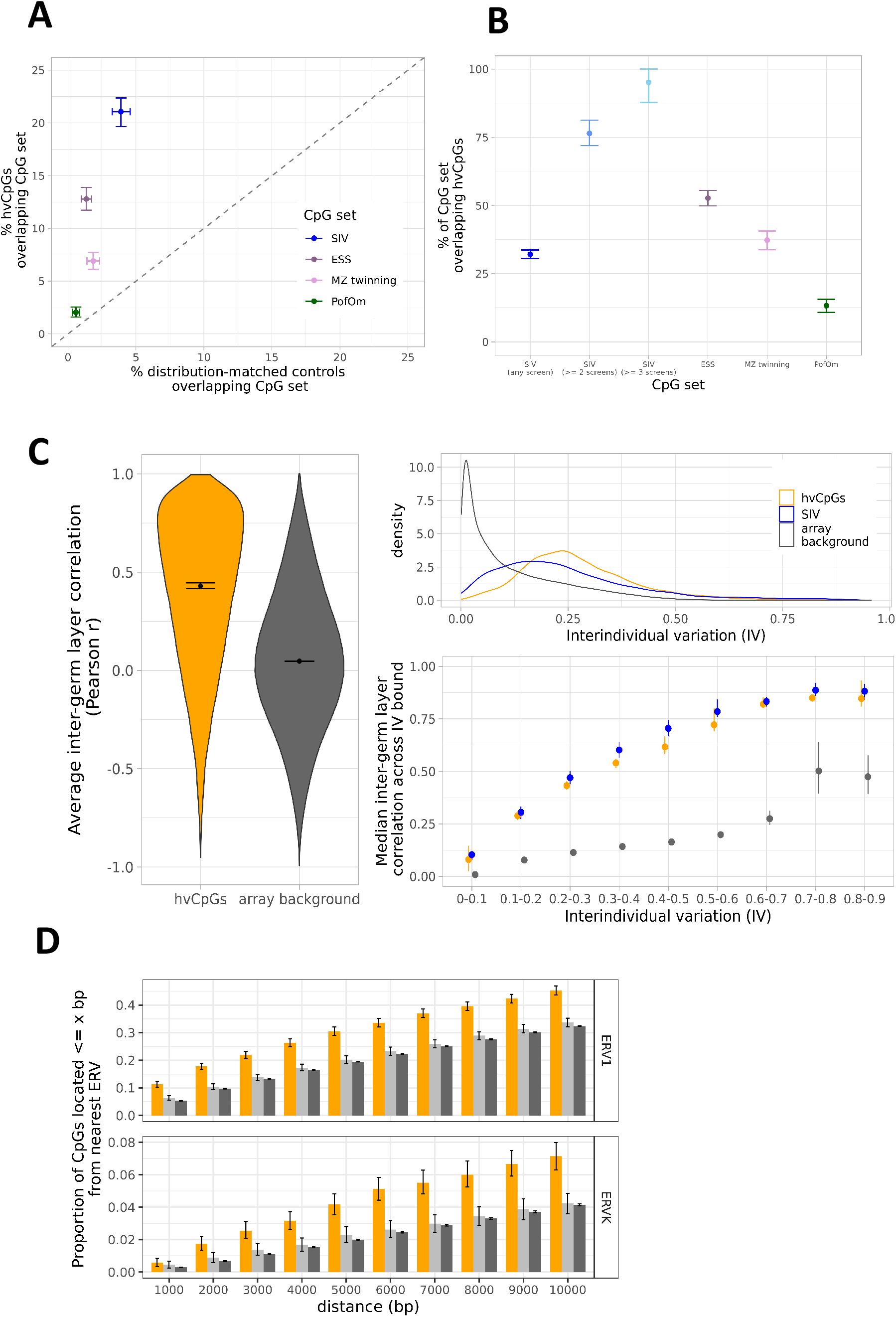
hvCpGs are enriched for loci and genomic features linked to variable methylation establishment in early development. **A)** The proportion of 3,566 hvCpGs (y-axis) vs corresponding distribution-matched controls (x-axis) covered in the ‘Blood_Cauc’ dataset that overlap 3,089 SIV-CpGs, 1,217 ESS CpGs identified by van Baak *et al*. (2018), 728 ‘MZ twinning’ CpGs identified by van Dongen *et al*. (2021) and 732 PofOm CpGs identified by Zink *et al*. (2018). **B)** The proportion of SIV-CpGs, ESS CpGs, MZ twinning CpGs and PofOm CpGs that are hvCpGs. SIV-CpGs identified in at least two or three independent screens were also included in this plot. **C)** Inter-germ layer correlations at hvCpGs using a fetal multi-tissue dataset that comprises methylation data from 10 individuals with endoderm- and ectoderm-derived tissues, 9 individuals with endoderm-and mesoderm-derived tissues and 8 individuals with mesoderm- and ectoderm-derived tissues (see Supplementary Table 7). **Left**: The distribution of average inter-germ layer correlations at 3,878 hvCpGs (orange) and 372,571 array background CpGs (excluding previously identified SIV CpGs and hvCpGs) (dark grey) covered in the fetal multi-tissue dataset. **Top Right**: Interindividual variation at 3,878 hvCpGs (orange), 4,076 previously identified SIV loci (blue) covered in the fetal multi-tissue dataset, and 372,571 array background CpGs (see ‘Methods’ for definition of interindividual variation). **Bottom Right:** Comparison of average inter-germ layer correlations at hvCpGs, SIV-CpGs and array background CpGs, stratified by interindividual variation. Each point indicates the median average inter-germ layer correlation for those CpGs with interindividual variation falling within each bound specified on the x-axis. **C)** The proportion of 3,566 hvCpGs, distribution-matched controls and array background CpGs that are ≤ *x* bp from the nearest ERV1 and ERVK transposable elements determined by RepeatMasker. Error bars in all panels are bootstrapped 95% confidence intervals. SIV = systemic interindividual variation, ESS = epigenetic supersimilarity, PofOm = parent-of-origin-specific methylation.

The set of all hvCpGs comprises 32.1% of the 3,089 CpGs reported as SIV in any of the four independent studies analysed despite comprising <1% of the 450K array. When considering ‘high-confidence’ SIV-CpGs reported in at least two or three of the four screens, the proportion identified rises to 76.5% and 95.1% respectively (Fig.3B). This suggests that our approach of identifying hypervariable loci across multiple datasets may be a more powerful method for identifying putative SIV loci, compared to existing SIV screens that necessarily rely on rare datasets with multi-tissue, mult*i*-germ layer methylation data from small numbers of individuals. To confirm this, we estimated the power to detect SIV using the multi-tissue data from four individuals analysed by van Baak *et al*.^25^. Using a permutation framework (‘Methods’), we estimated the mean power to detect SIV as 56% (median [IQR] = 0.58 [0.44, 0.72]; Supplementary Fig. 7). As expected, given the small sample size of this multi-tissue dataset, a large proportion of hvCpGs (75%) did not meet the minimum interindividual variation threshold of 0.2 used by van Baak *et al*. to define SIV. On the assumption that hvCpGs are highly enriched for true SIV, this could explain why hvCpGs constitute 61.7% of the van Baak *et al*. SIV-CpGs, while just 13.5% of hvCpGs are identified as SIV-CpGs in the van Baak *et al*. analysis.

To directly test our hypothesis that hvCpGs comprise previously unidentified SIV loci, we analysed a dataset of fetal tissues from 27 individuals, each with methylation data from two tissues derived from different germ layers (see Supplementary Table 7). Inter-germ layer correlations at hvCpGs had a median average Pearson r of 0.42, compared to array background CpGs which had a median average Pearson r of 0.05 (Fig. 3C left). Of the 3,878 hvCpGs covered in this fetal multi-tissue dataset, 1,653 (42%) had an average inter-germ layer Pearson r ≥0.5. Of these, 58% did not overlap previously identified SIV loci, suggesting that hvCpGs comprise novel SIV loci. A comparison of the average inter-germ layer correlation at hvCpGs and at previously identified SIV-CpGs showed that hvCpGs and SIV-CpGs had similar inter-germ layer correlations (Fig. 3C right).

### hvCpGs are enriched for loci with distinctive methylation patterns in MZ twins

We further investigated evidence for establishment of hvCpG methylation states in the early embryo by testing the overlap between hvCpGs and 1,217 “epigenetic supersimilarity” (ESS) CpGs overlapping array background. ESS CpGs show high interindividual variation with greater-than-expected methylation concordance between monozygotic co-twins in adipose tissue, suggestive of methylation establishment in the early zygote before MZ cleavage^25^. 13% of hvCpGs overlap an ESS CpG, showing a ∼9.5-fold enrichment for ESS CpGs relative to distribution-matched controls (Fig. 3A, Supplementary Table 12, Supplementary Fig. 10B).

We next examined the overlap between hvCpGs and a set of CpGs showing a unique methylation signature in adult tissues from MZ vs DZ twins (‘MZ twinning CpGs’, Table 2), implicating these CpGs in MZ twin splitting events in early development^28^. 7% of hvCpGs overlap an MZ twinning CpG, showing a 3.7-fold enrichment for MZ twinning CpGs compared to distribution-matched controls (Fig. 3A, Supplementary Table 12).

Notably, 54% of ESS and 37% of MZ twinning CpGs overlapping array background are hvCpGs (Fig. 3B).

### Reconciling the timing of variable methylation establishment at hvCpGs

The enrichments that we observe for SIV, ESS, evCpGs and MZ twinning CpGs offer a potential insight into the timing of methylation establishment at hvCpGs. 38% of hvCpGs overlap at least one of these CpG sets and enrichment is stronger amongst CpGs that show at least two of these properties (Supplementary Fig. 11). In particular, hvCpGs comprise 78% of SIV-ESS loci and 65% of SIV-MZ twinning loci, suggesting that SIV loci with evidence of establishment in the pre-gastrulation embryo are enriched for hvCpGs^26^.

Variable methylation states identified at evCpGs are thought to originate in embryonic development and/or early post-natal life^43^. We note that 41 out of 317 evCpGs overlap SIV and/or MZ twinning CpGs, suggesting that at least a subset may be established in the pre-gastrulation embryo. hvCpGs comprise 67% of evCpGs that overlap SIV-CpGs, and 76% of that overlap MZ twinning CpGs (Supplementary Fig. 11).

### hvCpGs are enriched for parent-of-origin methylation and proximal TEs

In mice, variable methylation states have been associated with the Intracisternal A Particle (IAP) class of endogenous retrovirus^58,59^, with growing evidence that methylation variability may in part be driven by incomplete silencing of IAPs in early development^60–62^. In humans, SIV-CpGs are enriched for proximal endogenous retrovirus elements (ERVs)^63^, including the subclasses ERV1 and ERVK^26^. This is also the case with hvCpGs, which show a ∼1.3-fold and ∼1.7-fold enrichment for proximal (within 10 kb) ERV1 and ERVK elements respectively, relative to both array background and blood distribution-matched controls (Fig. 3D, Supplementary Fig. 10C, Supplementary Table 12). Approximately 4.7% of hvCpGs are also located within 1Mb of telomeric regions, showing a 1.8-fold enrichment relative to distribution-matched controls and array background CpGs (Supplementary Table 12).

Maintenance of parent of origin-specific methylation (PofOm) in the pre-implantation embryo is critical for genomic imprinting^64^, and several previously identified SIV loci have been found to be associated with imprinted genes and/or PofOm^25,63^. 58 hvCpGs (1.4%) were annotated to 32 imprinted genes (Supplementary Table 13), no more than expected by chance since 1.9% of array background CpGs are annotated to imprinted genes. 10 hvCpGs were annotated to the polymorphically imprinted non-coding RNA *VTRNA2-1*, a well-established SIV locus that is associated with periconceptional environmental exposures^25,63,65–67^. Although only a small proportion (2.2%) of hvCpGs overlap regions of PofOm identified in peripheral blood^68^, this overlap represents a 3.5-fold and 11-fold enrichment relative to distribution-matched controls and array background respectively that is maintained after de-clustering (Fig. 3A, Supplementary Fig. 10D, Supplementary Table 12). This overlap constitutes 13% of all PofOm CpGs overlapping array background (Fig 3B).

### hvCpGs show sensitivity to pre-natal environment

Variable methylation states established in early development that are sensitive to environmental perturbation are promising candidates for exploring the developmental origins of health and disease^69–71^. We explored whether hvCpGs show sensitivity to pre-natal environment by examining their overlap with loci associated with season of conception (‘SoC’) in a rural Gambian population exposed to seasonal fluctuations in diet and other factors^72–74^. hvCpGs comprise 70 (29%) out of 242 previously identified SoC-CpGs^29^ overlapping array background, an approximately 3-fold enrichment relative to distribution-matched controls (Supplementary Table 11).

We next leveraged a recent meta-analysis of 2,365 cord blood samples that modelled genetic (G), genetic by environment (GxE) and additive genetic and environment (G+E) effects at variably methylated probes, where E represents a range of prenatal exposures including pre-pregnancy BMI, maternal smoking, gestational age, hypertension, anxiety and depression^14^. Of the 703 hvCpGs overlapping the neonatal blood variably methylated regions explored in that study, G, GxE, and G+E effects were the ‘winning’ models for 30%, 30% and 40% of probes respectively, representing an increase in G+E effects compared to array background (Supplementary Fig. 12). This analysis supports our intuition that hvCpGs are influenced but not determined by genetic variation, with pre-natal environment as an additional influencing factor.

### Chromatin states at hvCpGs

Compared to array background, hvCpGs are enriched within intergenic regions and CpG island ‘shores’ but are depleted within gene bodies and regions directly upstream of transcription start sties (Supplementary Fig. 9). We predicted chromatin states at hvCpGs by examining the overlaps of hvCpGs with histone modifications using the chromHMM 15-state model^52^ for seven tissues including embryonic stem cells (H1 ESCs), and fetal and adult tissues^53^. Although many hvCpGs were associated with regulatory elements in all tissues, hvCpGs were generally depleted in these regions compared to array background, except within predicted enhancers in H1 ESCs (Supplementary Fig. 13).

### Association with the clustered protocadherin gene locus on chromosome 5

Gene ontology enrichment analysis revealed that hvCpGs were significantly enriched for terms associated with cell-cell adhesion (Fig. 4A), which is largely driven by the colocalization of 3.3% of hvCpGs to clustered protocadherin (*cPCDH*) genes on chromosome 5. This region comprises three clusters of protocadherin genes (*cPCDHα, cPCDHβ, cPCDHγ*), each containing many variable exons whose promoter choice is determined stochastically via differential methylation by DNA-methyltransferase 3 beta (DNMT3B) in early embryonic development^75,76^, resulting in the expression of distinct *cPCDH* isoforms of cell-surface proteins that are critical for establishing neuronal circuits^77^. The *cPCDH* gene locus has also been found to be influenced by age^11,78–80^. Accordingly, although a minority (5%) of hvCpGs showed evidence of epigenetic drift in blood^11^, these are enriched within the *cPCDH* locus relative to those that did not show evidence of epigenetic drift (Fisher’s Exact Test (FET) p-value = 9.4 × 10^−9^, OR = 4.02). Hypervariable methylation states at the *cPCDH* gene locus may therefore be driven by early developmental and/or aging effects. Noting that evCpGs and MZ twinning CpGs (Table 2) have also been reported to colocalise with this locus^45,81^, hvCpGs annotated to *cPCDH* genes were ∼8.5-fold enriched for MZ twinning CpGs (FET p-value = 1.04 × 10^−22^) and ∼3-fold enriched for evCpGs (FET p-value = 1.6 × 10^−3^) relative to hvCpGs that were not.

**Figure 4.**
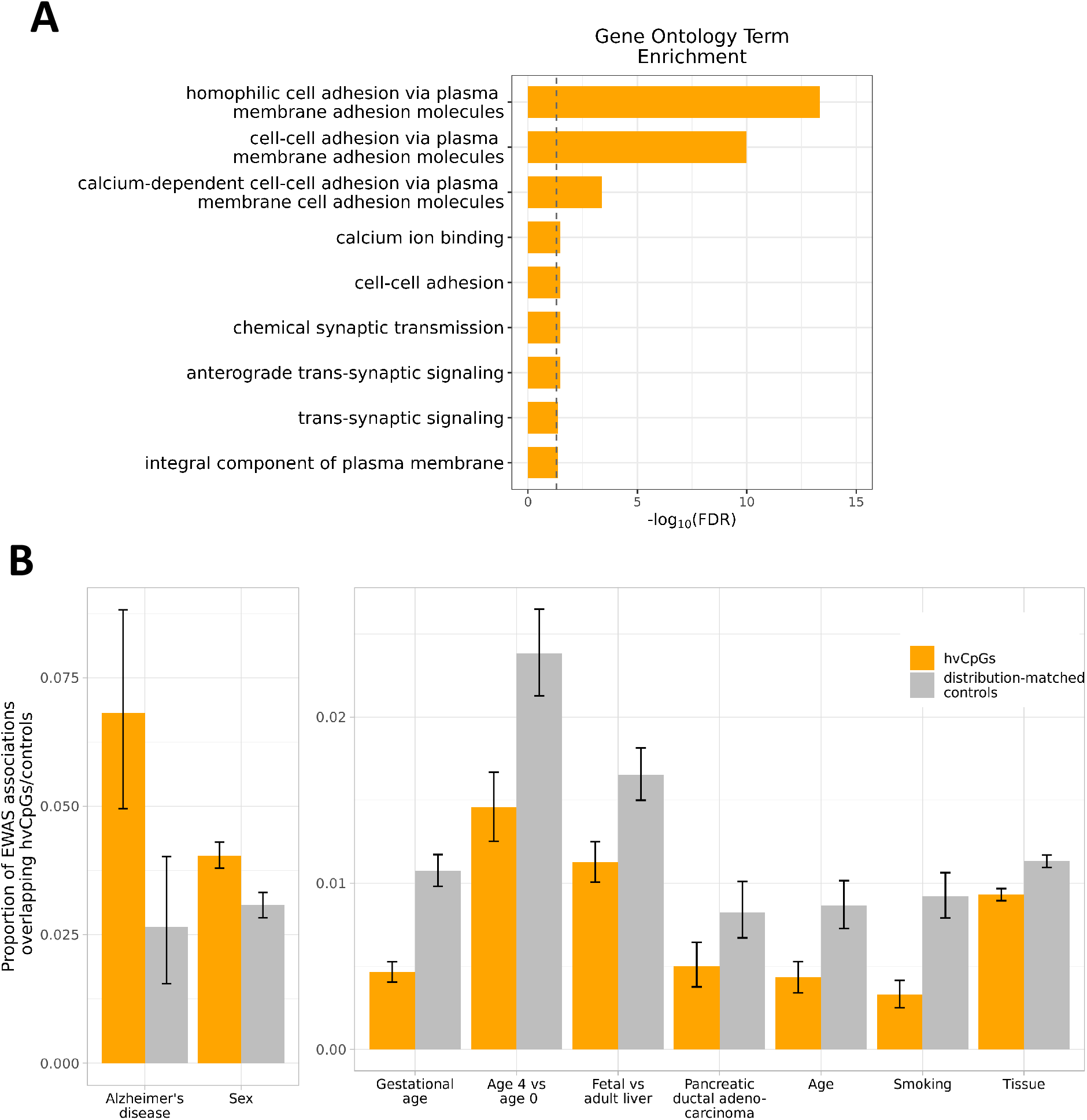
Functional annotation of hvCpGs. **A)** Gene ontology term enrichment analysis at hvCpGs. Vertical line indicates a significance threshold of FDR < 0.05. **B)** Enrichment (left) and depletion (right) of hvCpGs amongst EWAS trait associations relative to blood distribution-matched controls. Y-axis gives the proportion of EWAS trait associations that comprise hvCpGs and controls. Only traits overlapping at least 1% of hvCpGs were considered (see ‘Methods’ and Supplementary Table 9 for further details). Error bars are bootstrapped 95% confidence intervals.

### Association of hvCpGs with reported EWAS trait associations

To probe the potential functional role of hvCpGs, we analysed their overlap with traits reported in the epigenome-wide association studies (EWAS) catalogue (http://ewascatalog.org/). 86% of hvCpGs show significant associations (reported p-value < 1 × 10^−4^) with one or more of 231 unique traits covered in the catalogue (Supplementary Table 9). However, compared to blood distribution-matched controls, a suitable comparator given that the majority of EWAS have been carried out in blood, we found that hvCpGs were enriched amongst CpGs associated with sex and Alzheimer’s disease only (Fig. 4B, left panel).

Noting that sex-associated hvCpGs are not influenced by sex differences in the datasets that we analysed (Supplementary Fig. 6C, bottom) and that a similar proportion of SIV-CpGs are also associated with sex (23% of hvCpGs and 20% of the 3,089 SIV-CpGs considered in our study), we speculate that the association with sex may be a feature of variable methylation states established in early development. Amongst the 64 hvCpGs associated with Alzheimer’s disease, 23 overlap previously identified SIV and/or ESS loci, 9 of which annotated to *CYP2E1*, a gene that has also been associated with Parkinson’s disease and rheumatoid arthritis^82,83^.

hvCpGs were notably depleted amongst age-related traits relative to distribution-matched controls (Fig. 4B, right panel), in agreement with our earlier findings that hvCpGs are largely stable with age (Supplementary Fig. 6). hvCpGs are also depleted amongst CpGs that are differentially methylated between buccal cells and peripheral blood mononucleocytes (‘Tissue’ in Fig. 4B), supporting the notion that hvCpGs may be established before cell differentiation and that the method used to identify the hvCpGs is robust to tissue-specific methylation variation.

## Discussion

We have identified and characterised tissue- and ethnicity-independent hypervariable methylation states at CpGs covered by the 450k array. Our methodological approach was designed to be robust to dataset-specific drivers of methylation variability, including sex, age, cell type heterogeneity and technical artefacts. We identified 4,143 hvCpGs and found strong evidence that methylation states at many hvCpGs are likely to be established in the early embryo and are stable postnatally. Our analysis positions hvCpGs as tissue- and ethnicity-independent age-stable biomarkers of early stochastic and/or environmental effects on DNA methylation.

hvCpGs cover ∼1% of the 450K array and were in the top 5% variable methylation states in an average of 13 distinct tissues and 7 ethnicities. Our study is not the first to investigate DNAm patterns in multiple tissues. Previous studies have identified CpGs that are differentially methylated between tissues^85–87^; determined the extent to which variable methylation states in accessible tissues (such as blood) reflect those in inaccessible tissues such as brain^56,86–90^; compared methylation patterns between peripheral tissues^57,91,92^; directly identified SIV loci using tissues derived from different germ layers^24–27,63,93^; functionally characterised tissue-specific variably methylated regions^94^; and examined the extent to which common drivers of methylation variation, such as genetics, age, sex and environment, are tissue-specific^8,12,15–18,95,96^. The majority of these studies used a comparatively small number of tissues or cell-types, and few have used multi-tissue datasets from different ethnicities^15^.To our knowledge, ours is the first study to explore the extent to which variably methylated CpGs are shared across diverse tissues and ethnicities in the human genome.

The majority of hvCpGs were associated with at least one mQTL suggesting that additive genetic effects influence methylation variation at these loci. However, a comparison with mQTL-matched controls, together with evidence of enrichment for sensitivity to periconceptional environment and methylation discordance between MZ twins, suggests that stochastic and/or environmental effects have a relatively large influence on methylation variability at hvCpGs. In line with this, a large proportion of hvCpGs show evidence of systemic interindividual variation (SIV), that is, intra-individual correlation in methylation across tissues derived from different germ layers. Whilst loci that covary across different tissue types are enriched for mQTL effects^16,56,57,91^, it has been suggested that SIV loci are putative human metastable epialleles with variable methylation states established before gastrulation that are influenced but not determined by genetic variation^26^.

Our fetal multi-tissue analysis supports the notion that SIV at hvCpGs arises during development and is not, for example, driven by post-natal environmental influences that act across many tissues. hvCpGs were also highly enriched for epigenetic supersimilarity loci and MZ twinning-associated CpGs, both of which have been linked to establishment of methylation in the cleavage stage pre-implantation embryo^25,28^. The degree of overlap between variably methylated regions in different cell types has also been linked to their common developmental origin^94^. If this pattern holds true, it follows that stochastic and/or environmentally influenced variably methylated loci that are shared across a large number of diverse tissues are likely to have originated before germ-layer differentiation. We note that it is possible that variable methylation variation at some hvCpGs is influenced by later gestational or post-natal environmental effects, acting in addition to or independently of early environmental effects across multiple tissues, as has been suggested at the *VTRNA2-1* locus in the context of folate supplementation in pregnancy^97^, maternal age at delivery^67^, and smoking^98^.

The association of hvCpGs with parent-of-origin-specific methylation and proximal ERV1 and ERVK elements is notable because these features have been linked to SIV-CpGs^26^. This suggests that genomic regions targeted by epigenetic silencing or maintenance mechanisms during early embryonic reprogramming may be enriched for stochastic and/or environmentally influenced methylation variation. For example, it has been suggested that regions of PofOm may be vulnerable to stochastic or environmentally-sensitive loss of methylation on the usually-methylated allele or gain of methylation on the usually-unmethylated allele at a later time-point, leading to interindividual methylation variation^29,64,99^. Similarly, certain IAP elements (a class of ERVK LTR retrotransposon) show methylation variation between isogenic mice^58,59^ that in several cases can be influenced by pre-natal environment^100–103^. Whilst transposable elements are usually silenced to prevent insertion events from damaging the genome, recent evidence suggests that methylation variability at IAP elements is partly driven by low-affinity binding of *trans-a*cting Krüppel-associated box (KRAB)-containing zinc finger proteins (KZFPs)^60^ and by sequence variation in KZFP-binding sites^60,104^. Whilst KZFPs are known to target TEs in humans^105,106^, the extent of their role in driving methylation variation is an ongoing area of research.

The large overlap between hvCpGs and ‘high confidence’ SIV-CpGs identified in at least two independent screens suggests that the identification of hvCpGs might constitute a high-powered method for detecting novel SIV loci. Supporting this, the largest SIV screen to date with 10 individuals was reported to be underpowered to detect the well-established SIV locus at the non-coding RNA gene *VTRNA2-1*^27^ (represented by 10 hvCpGs), and we found that a 4-individual multi-tissue dataset analysed by van Baak *et al*.^25^ had limited power to detect SIV loci. Another consideration is that SIV screens to date have used different sets of tissues. Since loci that covary between one pair of tissues do not necessarily covary between another pair^56^, the enrichment for high confidence SIV loci might reflect the fact that methylation states at hvCpGs covary across a large number of tissues. Importantly, our analysis of a fetal multi-tissue dataset offers a strong validation of previously unreported SIV at hvCpGs.

Our analysis of EWAS trait associations revealed a moderate enrichment for hvCpGs amongst CpGs associated with Alzheimer’s disease and SIV loci have been linked to this and other disease outcomes including autism, cancer and obesity ^26,63,93^. For example, 10 hvCpGs overlap the *PAX8* gene which is a known SIV locus. *PAX8* methylation measured in peripheral blood of Gambian 2-year olds was recently shown to be correlated with thyroid volume and hormone levels in the same children in mid-childhood, and the latter was associated with changes in body fat and bone mineral density^107^. This suggests that hvCpGs are interesting candidates for exploring how stochastic and/or environmentally influenced DNAm states established in early development might influence life-long health.

hvCpGs are variable in diverse ethnicities, raising the possibility that regions of hypervariable methylation may be a conserved feature in the human genome. Stochastic methylation patterns established in the early embryo that are sensitive to early environment and that are able to influence gene expression might mediate a *predictive-adaptive-response* mechanism that senses the pre-natal environment in order to prime the developing embryo to its post-natal environment^70,71^. This would require environmentally responsive variable methylation states to be genetically hardwired into the human genome, providing a means of rapid adaptation to changing local environments on a scale much faster than is attainable through Darwinian evolution^108^. Associations between genotype and methylation variance have been previously reported, for example at the putative metastable epiallele *PAX8*^107^, at the master regulator of genomic imprinting *ZFP57*^25^ and at several probes in the major histocompatibility complex (MHC) region associated with rheumatoid arthritis^109^. Interestingly, 4% of hvCpGs are located within the MHC, representing an enrichment relative to the array background (FET p-value = 2.7 × 10^−10^, OR = 1.7). Further analysis of genotype-methylation variance effects is required to determine if this region, which contains a large amount of sequence variation and is implicated in many immune-mediated diseases^110^, might contain other examples of genetically-driven phenotypic plasticity that is mediated by DNA methylation.

Whilst our method of adjusting for the first 10 PCs of variation may not have controlled for all technical artifacts within each dataset, if technical issues were to cause a random CpG to be in the top 5% of variance in one dataset, this CpG would be unlikely to be in the top 5% of variance across a majority of datasets. The consequence would therefore be a loss of power to identify hvCpGs rather than the identification of spurious hypervariability. This is supported by our sensitivity analysis with unadjusted methylation data (Supplementary Fig. 8). An exception would be if the probe were consistently unreliable. We tested this using reliability metrics derived from analysis of technical replicates and found no evidence that hvCpGs are driven by technically unreliable probes. However, we note that better adjustment for technical artifacts within datasets^111^ and the addition of further datasets would likely lead to the identification of more hvCpGs.

The vast majority of publicly available methylation datasets use the Illumina 450k array. Therefore, a major limitation of this study is that we were only able to analyse the small proportion of the methylome covered by this array, which has been found to miss a disproportionate amount of variable CpGs^27^. However, we note that our method for identifying hypervariable CpGs can easily be applied to whole methylome sequencing data which is becoming increasingly available.

Through the joint analysis of methylation data from multiple tissues, we have identified a large set of hypervariable loci on the 450K array that are present across multiple tissues and ethnicities. Comparisons with a diverse range of data sources reveal that stochastic and/or environmentally-responsive methylation states at these loci are likely to have been established in the early embryo and appear to be stable with age, making them interesting candidates for studying the developmental origins of life-long health and disease.

## Data availability

The large majority of datasets analysed in this paper are in the public domain. GEO accession numbers and/or further details are provided in Supplementary Tables. A small number of analysed datasets have restricted access. Requests to access these should be submitted to the corresponding authors in the first instance with researcher access requiring an application to the relevant institutional review boards.

## Acknowledgements

GoDMC meta-GWAS summary statistics were kindly provided by Jordana Bell. This data has now been published (Min et al, 2021).

