## Supplementary Figures for "Tissue- and ethnicity-independent hypervariable DNA methylation states show evidence of establishment in the early human embryo"

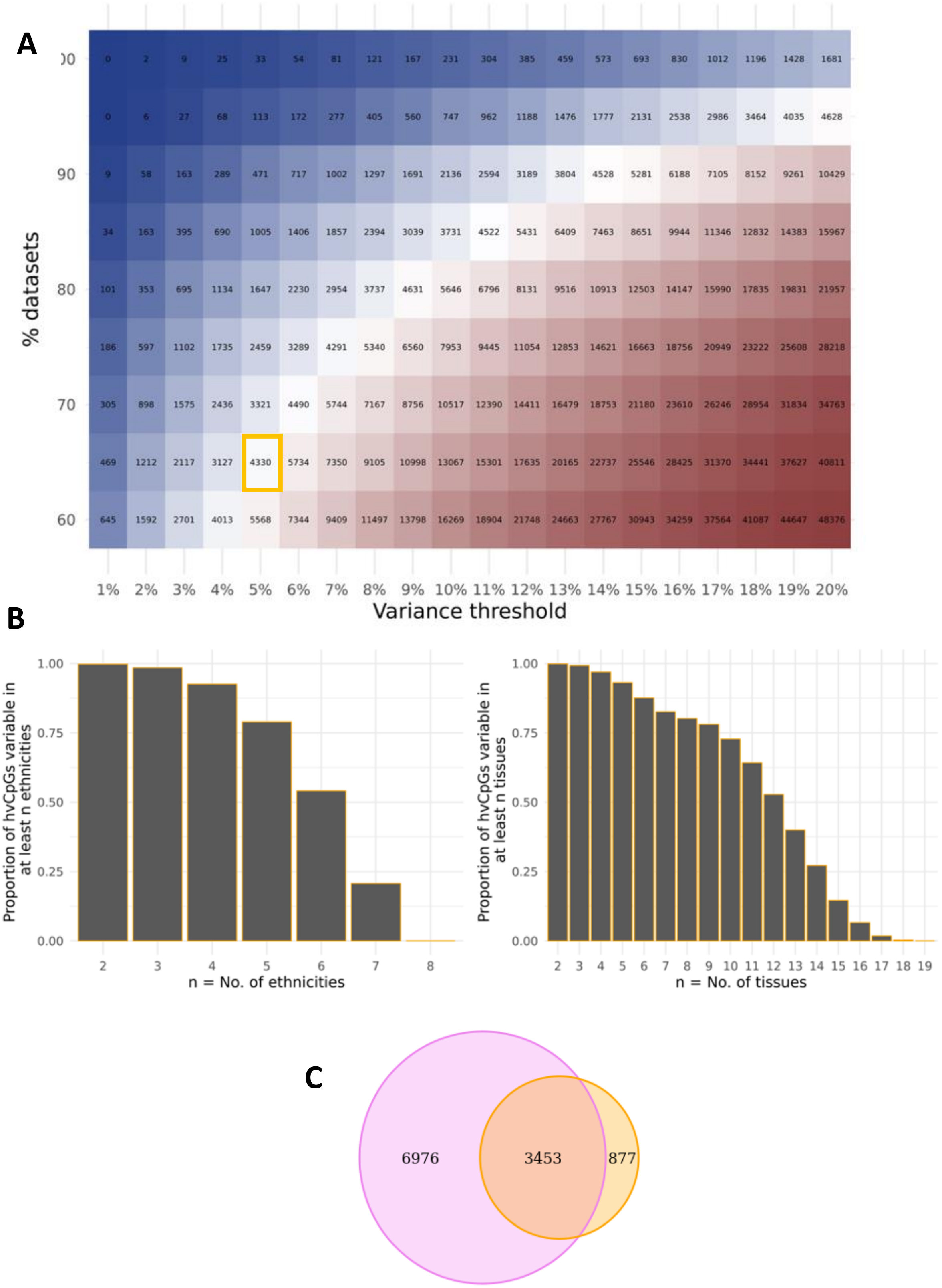


**Figure S1**. **Variable CpGs are common across diverse tissue types and ethnicities. A)** The number of CpGs within the top *i* % of variable sites by methylation Beta variance ('variance threshold') overlapping at least *j* % of datasets in which the CpG was present. CpGs were required to be present in at least 15 datasets. To identify hypervariable CpGs ('hvCpGs'), we set a threshold at *i,j* = *[5,65]*, marked by the orange box. **B)** The proportion of hvCpGs with top 5% beta variance in ≥ n ethnicities (left) and tissues (right). **C)** Overlap between the hvCpG set (yellow) and an alternative set obtained using *i,j* = [20,90] (purple).

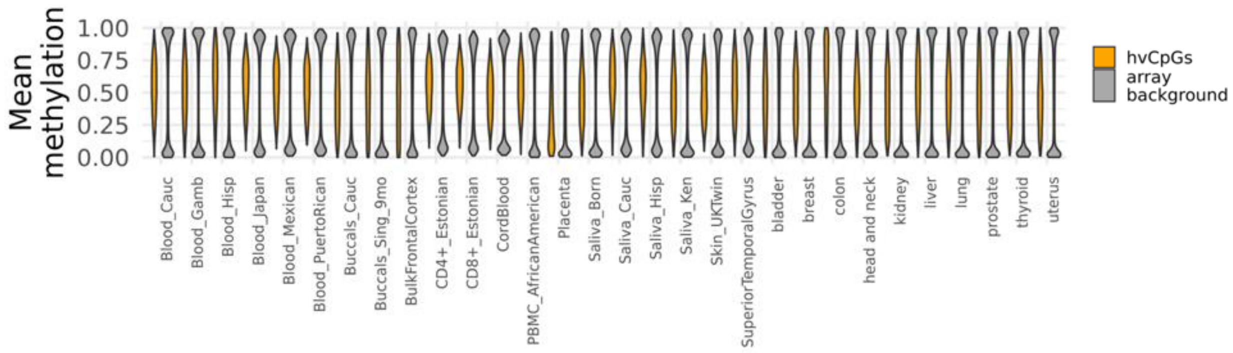


**Figure S2. hvCpGs are enriched for intermediate methylation values.** The distribution of mean methylation Beta values at hvCpGs (orange) and array background CpGs (grey).

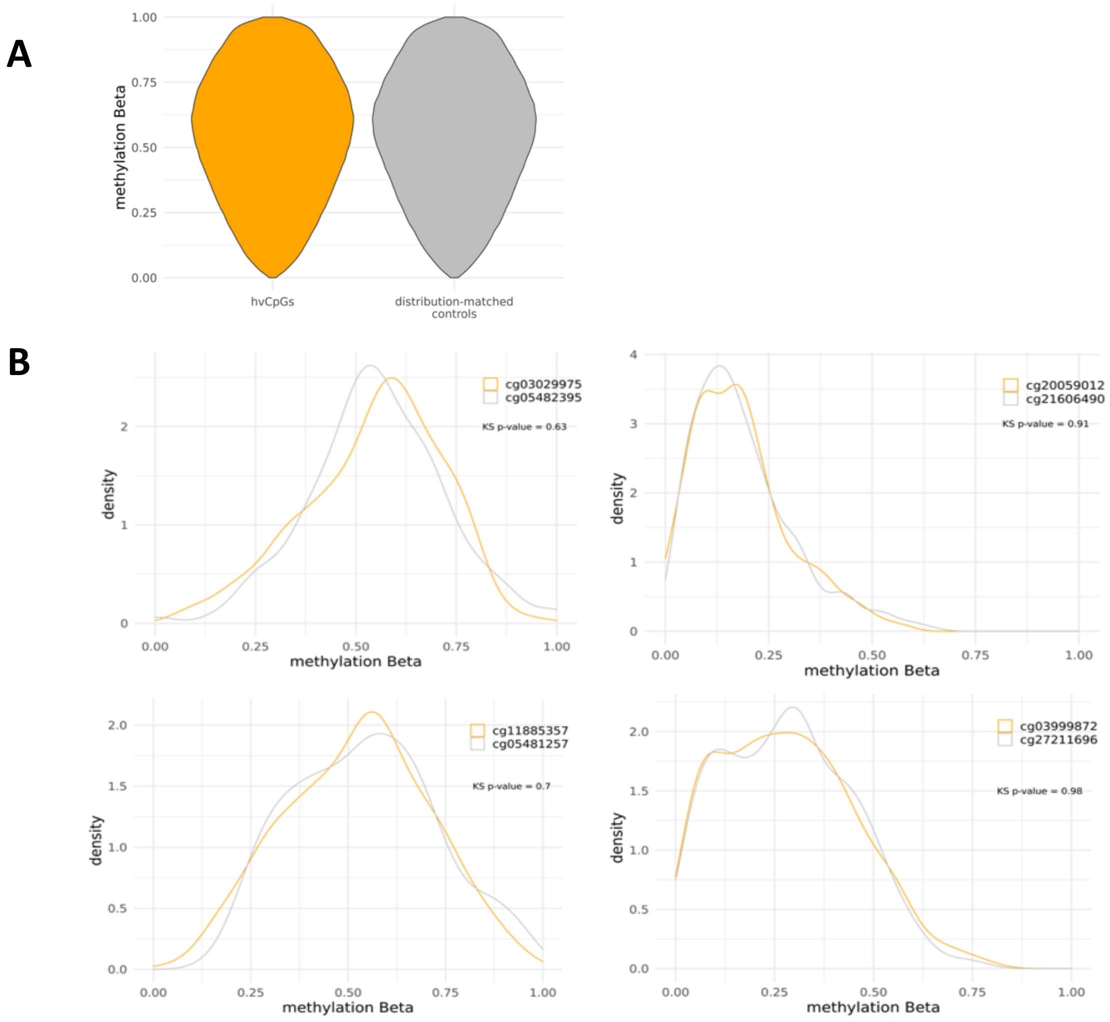


**Figure S3. Matching each hvCpG to a control with a similar distribution of methylation Beta values in Caucasian blood. A)** The distribution of methylation Beta values at 3,566 hvCpGs and their corresponding distribution-matched controls selected from the 'Blood_Cauc' dataset (Supplementary Table 1). **B)** Examples of the similar distribution of methylation Beta values at hvCpGs (orange) and their corresponding controls (grey). The median Kolmogorov-Smirnov p-value over all 3,566 hvCpG-control pairs was 0.91 (IQR = [0.63, 0.98]). See 'Methods' for further details.

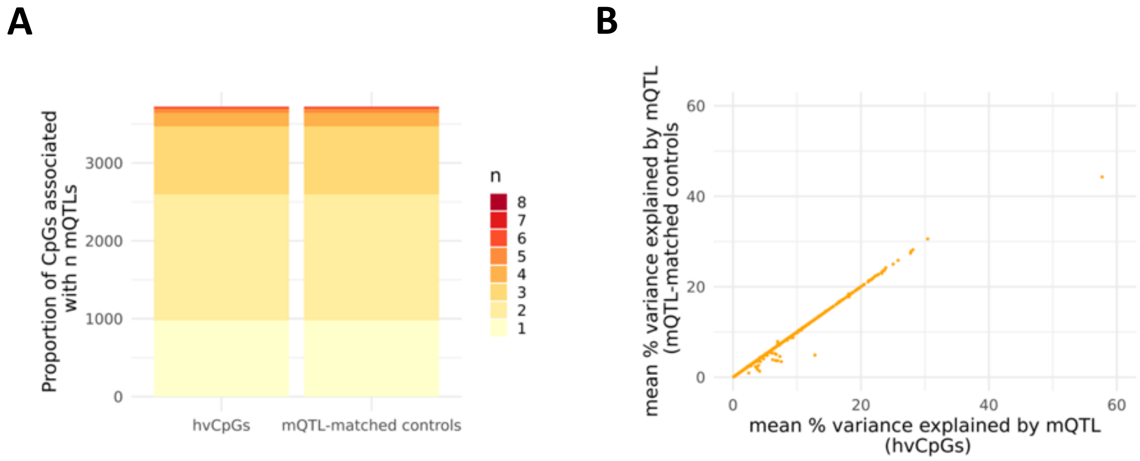


**Figure S4. Matching each hvCpG to a CpG with similar mQTL effects using the GoDMC mQTL analysis.** Each of the 3,722 hvCpGs reported in the GoDMC mQTL analysis (Min et al. 2021), was matched to a control CpG with **A)** the same number of mQTL associations, and **B)** similar mean % variance explained by mQTL, requiring each control CpG to be present in at least as many datasets as the hvCpG.

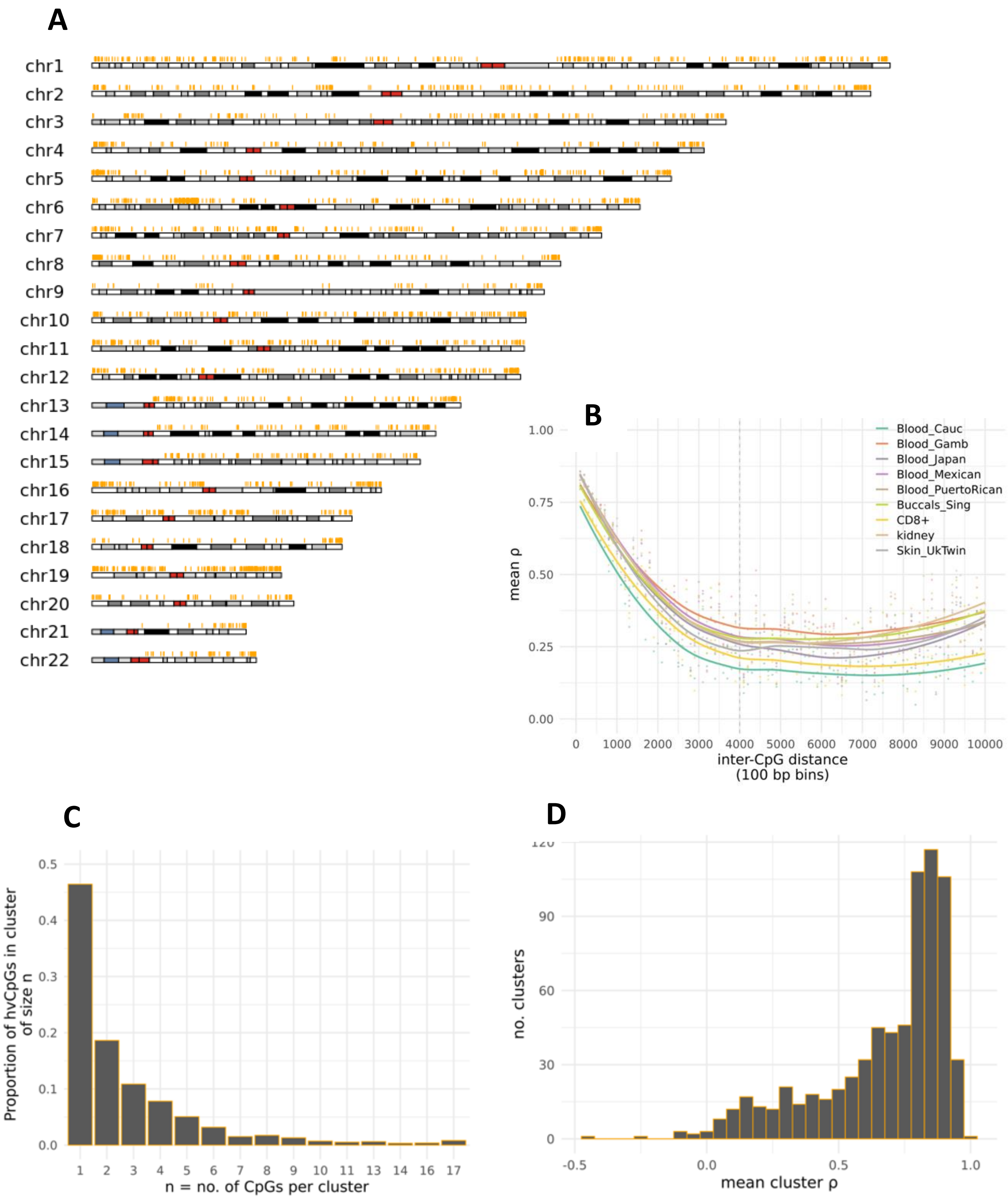


**Figure S5. Clustering of hvCpGs. A**) The positions of hvCpGs across autosomal hg19 chromosomes. **B)** The decay of methylation correlation with distance at hvCpGs. Each point indicates the average pairwise Spearman rho across hvCpG pairs with inter-CpG distance falling within each 100 bp bin. Curves are loess lines of best fit for each dataset. Dashed vertical line at 4000 bp is the threshold we used for defining hvCpG clusters. **C)** The proportion of hvCpGs that fall into clusters comprising *n* CpGs, using an inter-CpG distance of 4000 bp to define clusters. **D**) The average Spearman rho across the 716 hvCpG clusters that comprise at least 2 CpGs. For each cluster, we calculated the average pairwise Spearman rho across hvCpG pairs for every dataset in which all CpGs in the cluster were covered, before taking the mean across these datasets. Note, in A and C we only considered datasets with N ≥ 100.

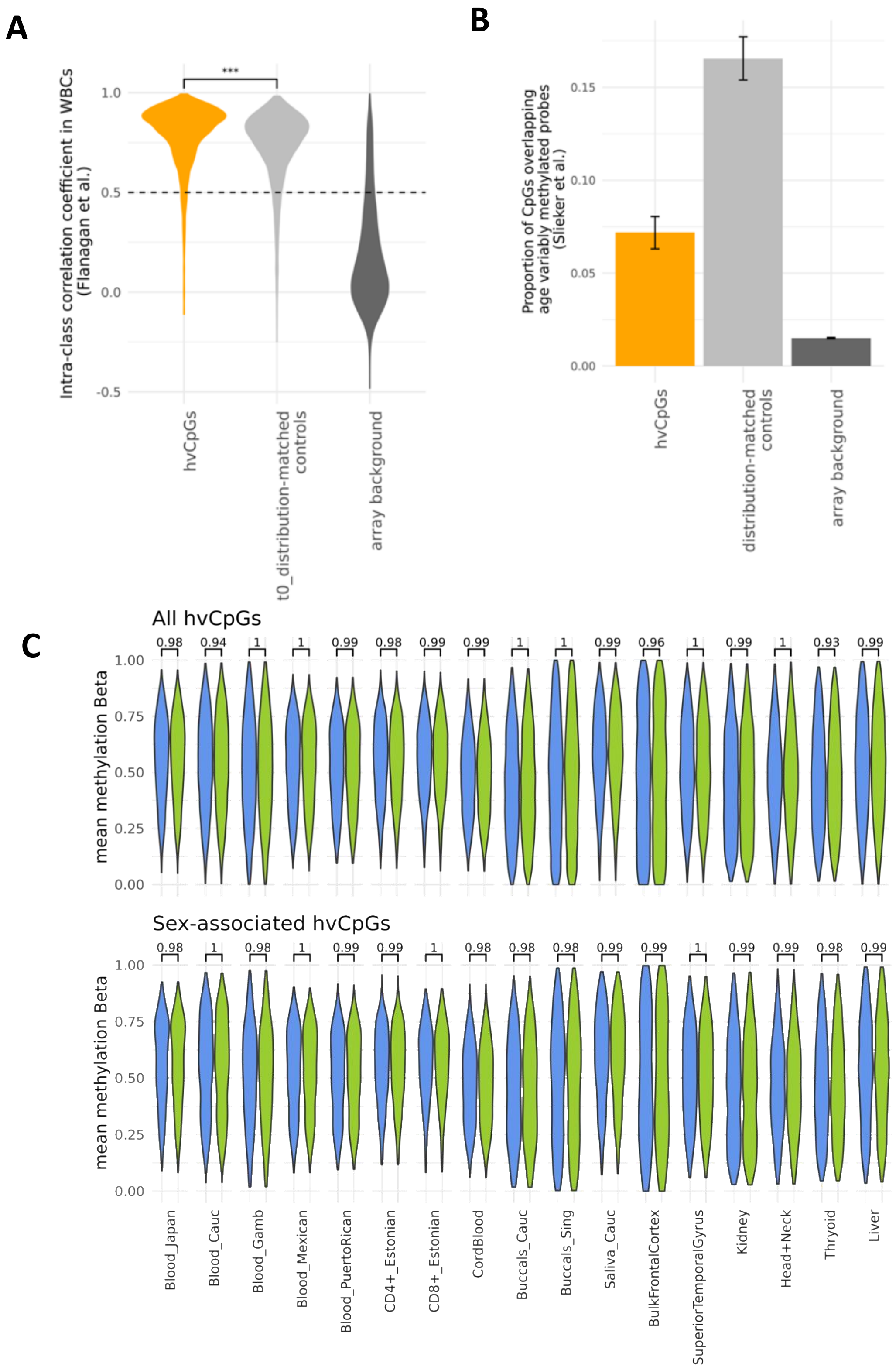


**Figure S6. Methylation at hvCpGs is stable with age and is not influenced by sex. A)** The distribution of temporal ICC scores at 3,564 hvCpGs and CpGs with similar distribution of methylation Beta values at the first time-point (t0) covered in the Flanagan et al. (2015) dataset (see Methods). The horizontal line at ICC = 0.5 indicates the threshold specified by Flanagan et al. (2015) above which probes are considered to be temporally stable. *** indicates Wilcoxon paired signed-rank test p-value < 0.001. **B)** The proportion of 3,566 hvCpGs and corresponding 'distribution-matched controls' (Table 1) covered in the 'Blood_Cauc' dataset that overlap probes showing increased variability with age in whole blood in a study of 18-to-88-year olds by Slieker et al.(2016). **C)** Comparison of the distribution of mean methylation Beta values between male (blue) and female (green) samples at all hvCpGs (top panel) and at 941 hvCpGs associated with sex in the EWAS catalog (bottom panel) for datasets with N $\geq$ 50**.** Wilcox Rank Sum p-values are given above each pair of violin plo

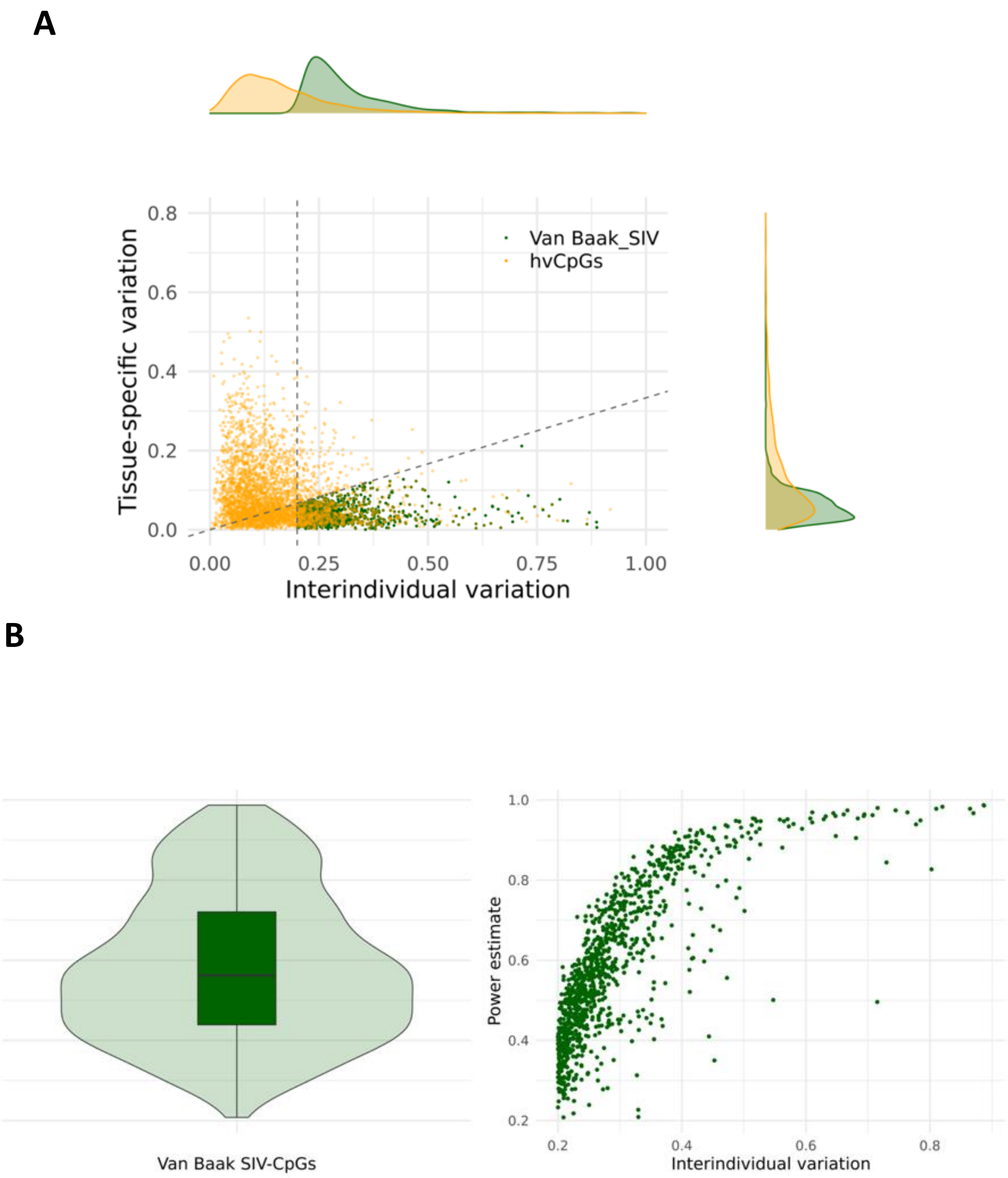


**Figure S7. Estimating the power to detect SIV-CpGs reported by van Baak et al. (2018) using a 4-individual multi-tissue dataset (GSE50192). A)** Replication of the screen by van Baak et al., in which 1,042 SIV-CpGs (green) were identified, using the same methylation data from abdominal aorta, gall bladder and ischiatic nerve in 4 individuals. Orange points are hvCpGs. **B**) Estimating the power to detect each of the 1,042 SIV-CpGs reported by van Baak et al. (2018). See Methods for further details**. Left:** distribution of power estimates across 1000 permutations**. Right:** confirmation that estimated power to detect is correlated with interindividual variation.

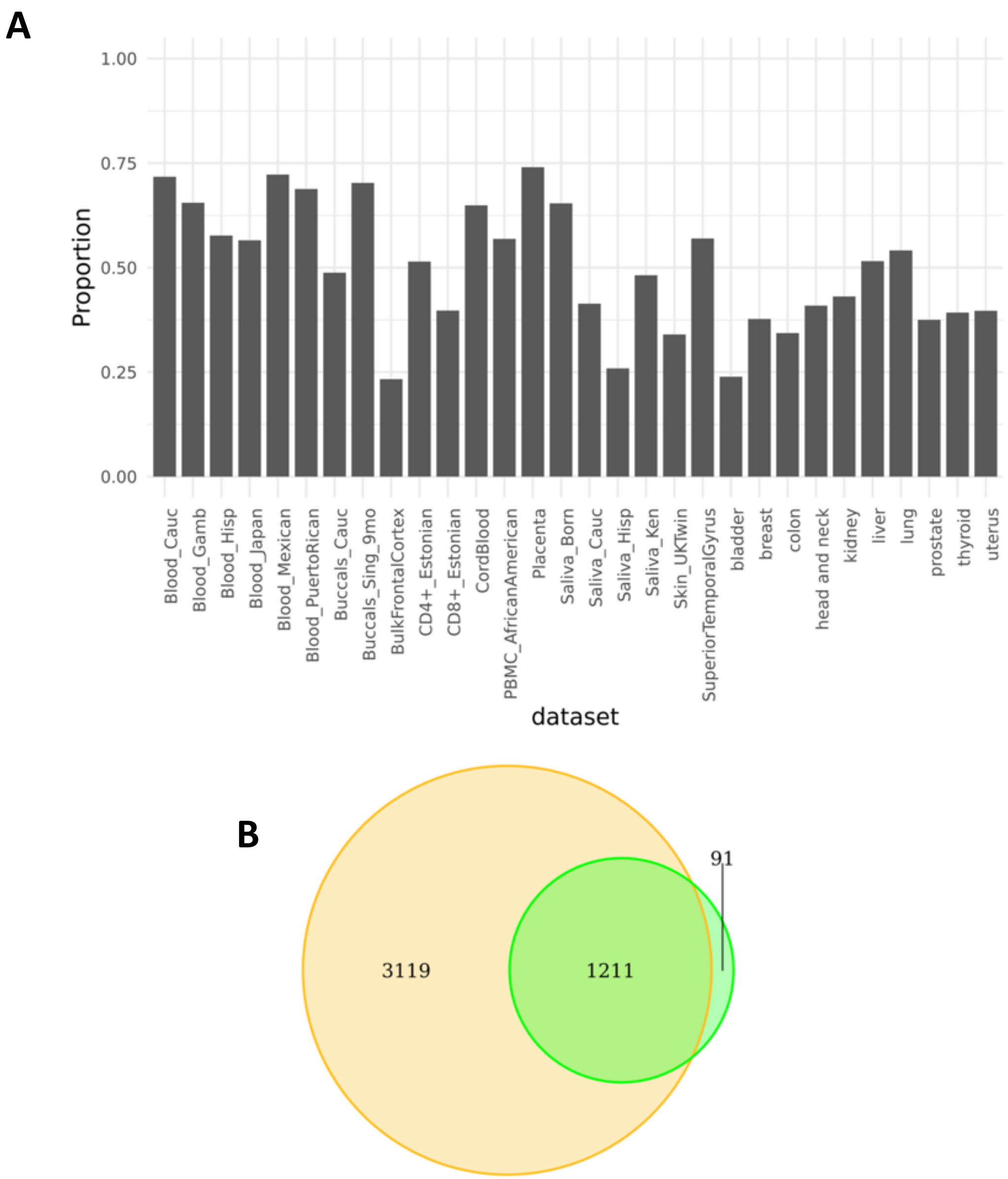


**Figure S8. Adjustment for the first 10 PCs of methylation variation in each dataset increases the number of hvCpGs. A)** For each dataset, we extracted the top 5% of CpGs by methylation Beta variance before and after adjusting methylation for the first 10 PCs. We then determined the proportionate overlap between PC-adjusted and PC-unadjusted variable CpGs. As expected, PC-adjustment influences the most variable CpGs within each dataset. **B)** The overlap between the set of 4,330 hvCpGs identified using PC-adjusted datasets (orange) and a set of 1,302 hvCpGs identified using unadjusted datasets (green). PC-adjusted: $lm(meth \sim 10\ PCs + age + sex)$, unadjusted: $lm(meth \sim age + sex)$.

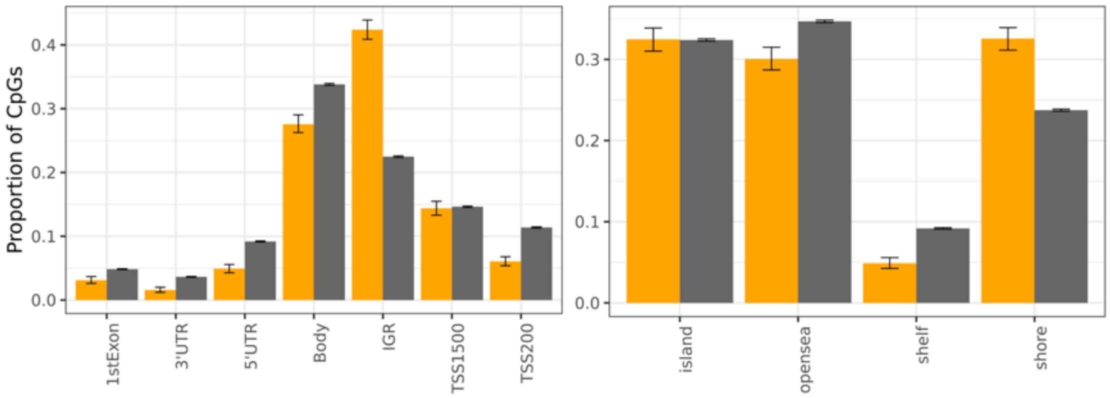


**Figure S9. Genomic locations of hvCpGs (orange) vs array background (grey) in relation to gene bodies (left) and CpG islands (right).** UTR = untranslated region, IGR = intergenic region, TSS = transcription start site. Error bars are bootstrapped 95% confidence intervals.

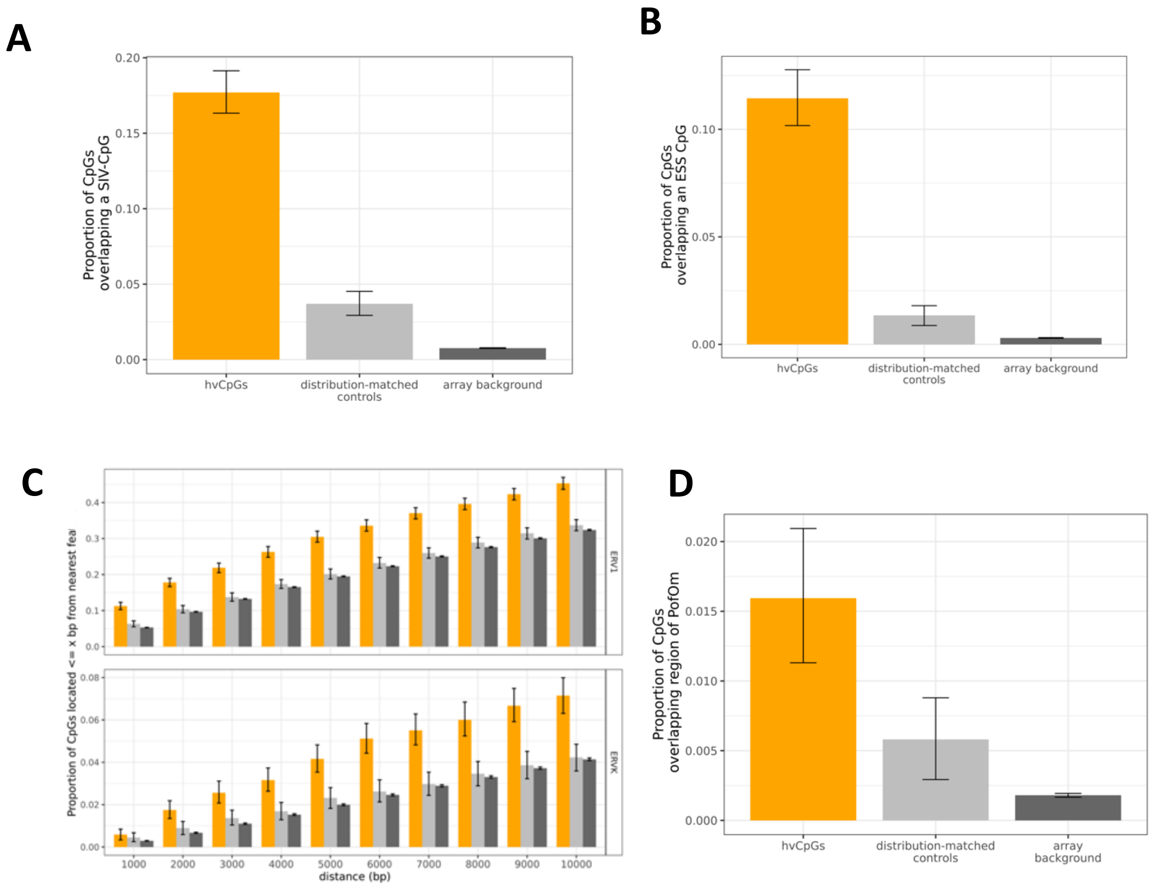


**Figure S10. The association between hvCpGs and features suggestive of methylation establishment in the early embryo is maintained when using a de-clustered set of hvCpGs.** The proportion of 2,178 de-clustered hvCpGs (in which no CpG is within 4 kb from another) and corresponding distribution-matched controls covered in the 'Blood_Cauc' dataset that **A)** overlap SIV-CpGs (4.8-fold enrichment relative to controls)**, B)** overlap ESS-CpGs (8.5-fold enrichment)**, C)** are located $\leq x$ bp from nearest ERV1 and ERVK transposable elements, and **D)** overlap regions of parent-of-origin specific methylation (PofOm) identified by Zink et al. (2018) (2.7-fold enrichment).

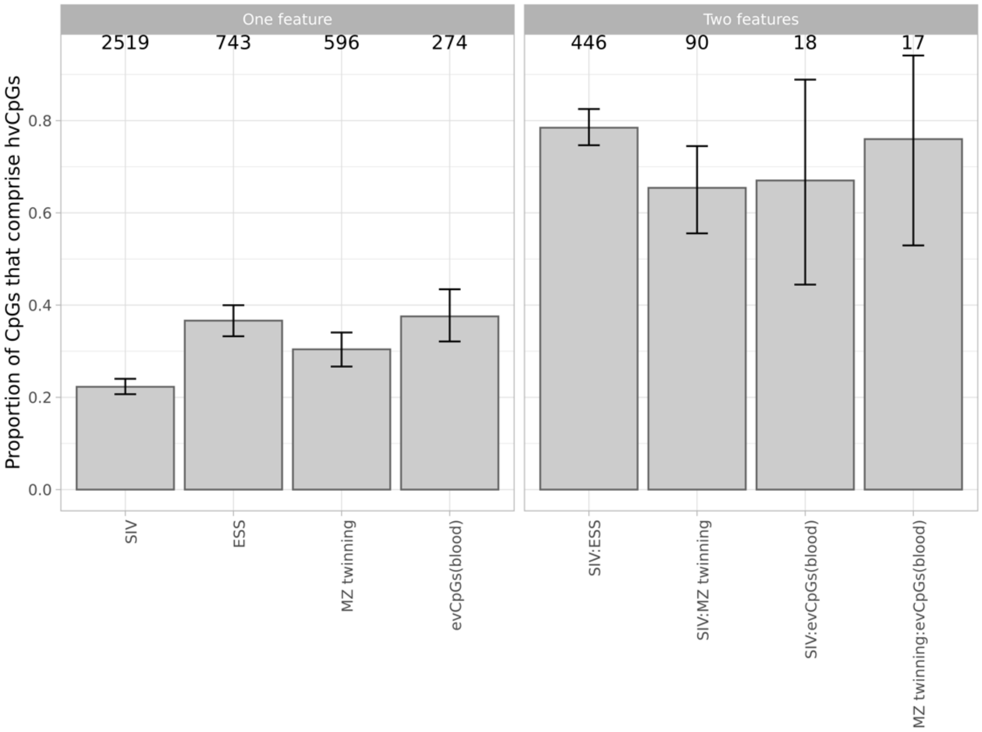


**Figure S11. hvCpGs comprise a large proportion of loci that show more than one feature linking them to variable methylation establishment in the early embryo.** Number of CpGs overlapping the array background is given above each bar, requiring a minimum of 15 CpGs. Error bars are bootstrapped confidence intervals. SIV = systemic interindividual variation, ESS = epigenetic supersimilarity, evCpGs = equivalently variable CpGs.

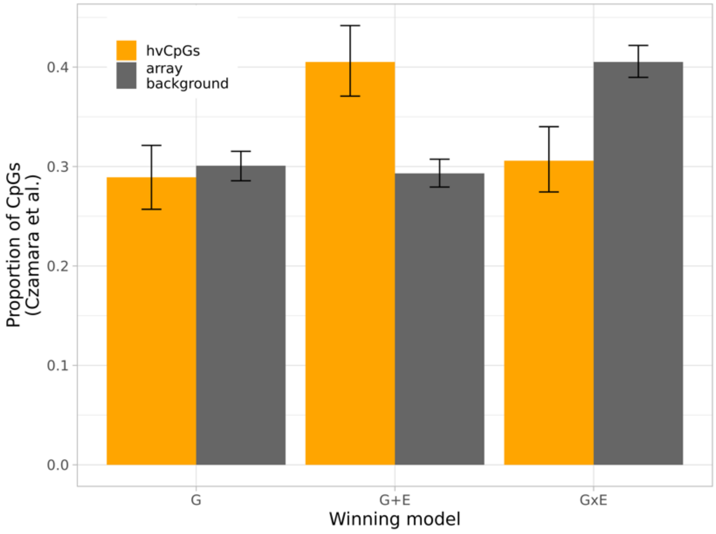


**Figure S12. Variability of hvCpGs in whole blood is best explained by G+E effects estimated by Czamara et al. (2019**). The proportion of 747 hvCpGs and 3,644 array background CpGs covered by the variably methylated probes (VMPs) reported by Czamara et al. that are best explained by G, G + E and G x E effects. Error bars are bootstrapped 95% confidence intervals.

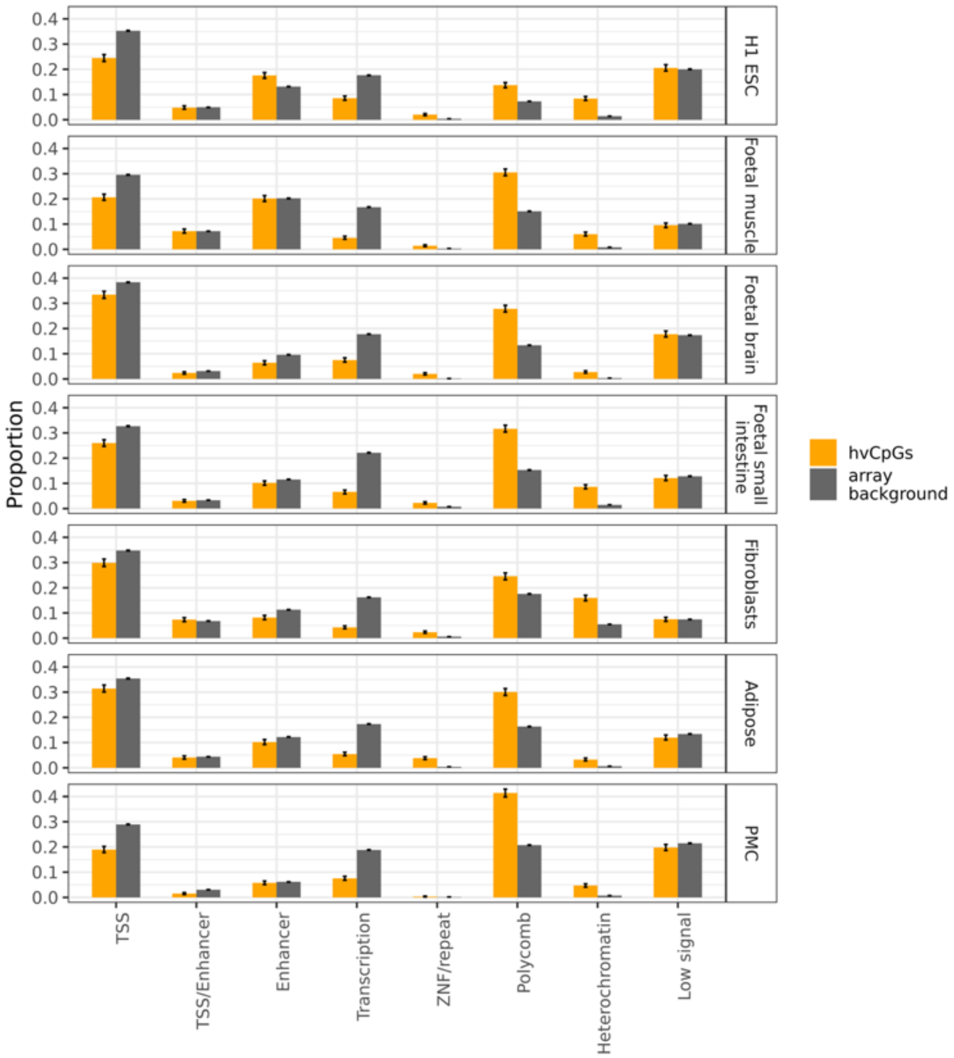


**Figure S13. Functional annotation of hvCpGs using the ChromHMM model.** The chromHMM 15-state model using data from H1ESC, foetal muscle, foetal brain, foetal small intestine, fibroblasts, adipose and primary mononucleocytes from the Roadmap Epigenomics Consortium. 15-states have been collapsed to 8 states for clarity (Supplementary Table 8). TSS = transcription start site. ZNF = zinc finger gene. Error bars indicate bootstrapped 95% confidence intervals.
